# Loss of intestinal ChREBP impairs absorption of dietary sugars and prevents glycemic excursion curves

**DOI:** 10.1101/2021.05.18.444615

**Authors:** W Charifi, V Fauveau, L Francese, A Grosfeld, M Le Gall, S Ourabah, S Ellero-Simatos, T Viel, M Cauzac, D Gueddouri, F Benhamed, B Tavitian, R Dentin, AF Burnol, C Postic, S Guilmeau

**Author notes:** Correspondence should be addressed to: Sandra Guilmeau Institut Cochin, Department Endocrinology Metabolism & Diabetes, 24 rue du Faubourg Saint Jacques, 75014 Paris, France.

## Abstract

Increased sugar consumption is a risk factor for features of the metabolic syndrome including obesity, hypertriglyceridemia, insulin resistance, diabetes, and nonalcoholic fatty liver disease. The gut epithelium, which plays a central role in dietary sugar digestion, absorption and metabolism has emerged a key actor of metabolic disorders. While the transcription factor ChREBP (Carbohydrate response element binding protein) has been established as a key player of the adaptive reprograming of cellular metabolism in various tissues upon glucose or fructose challenge, its specific contribution to the regulation of blood glucose upon dietary sugar intake was not previously addressed.

We demonstrate here that ChREBP is abundantly expressed in the proximal gut epithelium, where carbohydrate digestion and absorption primarily occur and in particular L cells, which produce the glucoincretin GLP-1. The inducible deletion of ChREBP specifically in the mouse gut epithelium (Ch^ΔGUT^ mice) resulted in the reduction of early glycemic excursion upon oral glucose load. Surprisingly, despite being associated with reduced GLP-1 production, loss of gut ChREBP activity significantly dampened glucose transepithelial flux, and thereby delayed glucose distribution to peripheral tissues. Among the underlying mechanisms, we unveil that Ch^ΔGUT^ mice show an impaired expression of key intestinal hexose (glucose, galactose, fructose) transporters and metabolic enzymes as well as brush border dissacharidases. In agreement, intestinal ChREBP deficiency was accompanied by a precocious intolerance to both high-lactose and high-sucrose diets concomitant with mild galactose and severe fructose malabsorption syndromes.

Altogether, our study demonstrates that, by transcriptionally orchestrating local digestion and absorption of dietary sugars, ChREBP activity in the mouse gut epithelium controls glucose appearance rate into systemic circulation and prevents against intolerance to mono- and disaccharides.

## INTRODUCTION

Prominent dietary sugars include sucrose and lactose, and their constituent monomers, glucose, fructose, and galactose. While the consumption of dietary sugars has tripled worldwide over the last five decades, mostly due to excessive intake of added fructose, global health guidelines are calling for reduction in consumption of dietary sugars due to concerns over their potential roles in disease onset. Epidemiological studies have documented a strong correlation of enhanced sugar intake with numerous pathological conditions associated with impaired glycemic control, such as obesity, diabetes, and non-alcoholic fatty liver disease (Jegatheesan & De Bandt, 2017a; Stanhope, 2012). In this context, the gut occupies a decisive position as it contributes to sweet tasting, to early breakdown and absorption of sugars as well as to incretin response and neural regulation of food intake, which are all key players in the regulation of postprandial glycemia. This is illustrated by blood glucose-lowering approaches of type 2 diabetes treatment targeting either the activity of enterocyte brush border alpha-glucosidases (acarbose, miglitol) (Chiasson et al., 2002; Hanefeld, 2007; Inzucchi et al., 2015; Yang et al., 2014) or the insulin secretagogue action of enteroendocrine L cells-derived GLP1 (GLP1R agonists, DPP4 inhibitors) (Gilbert & Pratley, 2020).

Interestingly, recent evidence supports that sugar metabolism represents another important feature of gut epithelial cell potency to control glycemia, in particular upon dietary or blood sugar overload. Thus, while the small intestine was recently identified as the major site of dietary fructose clearance in mice, it was proposed that its metabolic capacity to convert it into glucose determines liver exposure to portal fructose and thereby fructose hepatic toxicity (Jang et al., 2018, 2020). Moreover, gut alterations in intracellular glucose metabolism upon hyperglycemic conditions were reported to interfere with intestinal permeability, a critical player of metabolic low-grade inflammation (Thaiss et al., 2018). In addition, monosaccharide consumption has been shown to enhance intestinal *de novo* lipogenesis, and glucose or fructose were highlighted as substrates for triglyceride synthesis and lipoprotein export in the form of chylomicrons (Hoffman et al., 2019). Of note, the contribution of intestinal sugar metabolism to glycemic control was also reported upon fasting state, as intestinal glucose production is essential to maintain glucose homeostasis in the absence of hepatic glucose production under this condition (Penhoat et al., 2014).

In this context, the transcription factor ChREBP (Carbohydrate response element binding protein), which is encoded by the *mlxipl* gene, was demonstrated to mediate the transcriptional effects of glucose and fructose on genes encoding enzymes of key intracellular metabolic pathways (glycolysis, fructolysis, *de novo* lipogenesis) in the liver or in the white adipose tissue (Abdul-Wahed et al., 2017). Interestingly, whole-body knockout of *mlxipl* was primarily associated with a pronounced intolerance to high-sucrose challenge and to fructose malabsorption in mice (Iizuka et al., 2004; Oh A et al., 2018; Kato T et al., 2018). *Kim et al.* further demonstrated that intestinal rather than hepatic ChREBP activity is essential for fructose tolerance and that constitutive invalidation of ChREBP in the gut is sufficient to recapitulate the fructose-mediated toxicity observed in global ChREBP knockout mice (M. Kim et al., 2017). While intestinal epithelial cells are key players for the release of glucose into the portal circulation upon fructose or glucose challenge (Gorboulev et al., 2012; Jang et al., 2018; Schmitt et al., 2017), and reduced expression of hexose transporters was previously reported in the gut of mouse models with total ChREBP deficiency in response to fructose or sucrose (Kato et al., 2018; Oh et al., 2018), the specific contribution of intestinal ChREBP activity to the regulation of blood glucose upon dietary sugar intake was not previously addressed.

To test this hypothesis, we generated an inducible model of ChREBP invalidation in intestinal epithelial cells (Ch^ΔGUT^ mice). Interestingly, specific gut deletion of ChREBP in adult mice was associated with reduced glycemic excursion upon oral glucose or fructose load. At the mechanistic level, we unveiled that intestinal ChREBP activity controls glucose transepithelial flux and orchestrates the expression of intestinal hexose (glucose, galactose, fructose) transporters and metabolic enzymes as well as brush border dissacharidases. Consequently, Ch^ΔGUT^ mice displayed an early intolerance to both high-lactose and high-sucrose diets concomitant with mild galactose and severe fructose malabsorption syndromes. Altogether, our data pinpoint that intestinal ChREBP activity is a pivotal transcriptional mediator of the adaptative response to luminal dietary sugars in the gut epithelium and of the consecutive glucose appearance rate into systemic circulation.

## MATERIEL AND METHODS

### Generation of transgenic mice with targeted deletion of Mlxipl in the intestine

Heterozygous *mlxipl*^tm1a(EUCOMM)Wtsi^ mice were generated as previously described (Iroz et al., 2017), using a targeting vector with an hygromycin cassette that was inserted between exon 15 and exon 16 of the murine *mlxipl* gene, flanked by two FRT (Flipase Recognition Target) sites and two loxP sites flanked exon 9 and hygromycin cassette. These founder mice were crossed with mice that express the Flippase recombinase to remove the hygromycin cassette, thereby generating *mlxipl^fl/fl^* mice (Figure S1). *Ch^fl/fl^* mice were then crossed with *villin-Cre^ERT2^* mice, which specifically express the Cre recombinase in the gut epithelium (El Marjou et al., 2004) to generate *mlxipl^fl/fl;Vil-CreERT2^* mice. Intestinal inducible deletion of *mlxipl* (Ch^ΔGUT^) was performed in 8 week old *mlxipl^fl/fl^*^;Vil-CreERT2^ mice through daily gavage with tamoxifen citrate salt (1mg/mice) (Sigma T9262) diluted in 5% carboxy-methylcellulose sodium salt during 5 consecutive days. Similarly, *mlxipl^fl/fl^* mice were given tamoxifen and were used as controls (CTL mice). Mice were genotyped using the following PCR primers: *Villin-Cre* F: 5’ CAG-GGT-ATA-AGC-GTT-ATA-AGC-AAT-CCC 3′ and R: 5’ CCT-GGA-AAA-TGC-TTC-TGT-CCG 3′; *mlxipl floxed* F: 5’ CAC-TGA-GTG-TCC-ACC-TGT-CTC-CCC-CT 3′ and R1: 5’ GCA-CCC-ATT-TAC-CAA-CTT-AGT-C 3′ and R2: 5’ TCC-CAC-ATC-TCT-AGG-CTC-AG 3′.

### Animals and dietary challenges

Experimental procedures, which were conforming to the French guidelines for animal studies, were approved by the local Animal Care and Use Committee (CEEA Paris Descartes) and, were registered at the National Ethics Committee (CNREEA Ile-de-France no. 34, agreement # 12-129 and 12-162).

8- to 12-week-old C57BL/6J (Janvier Labs), Ch^−/−^ (Iizuka et al., 2004), Ch^ΔGUT^ mice and their respective control littermates (CTL) were used for all experiments. Ch^−/−^ and Ch^+/+^ mice as well as Ch^lox/lox^ and Ch^ΔGUT^ mice were co-housed and maintained in a 12-hr light/dark cycle with free access to water and standard diet (65% carbohydrate, 24% protein and 11% fat) unless otherwise specified. Mice Ch^lox/lox^ and Ch^ΔGUT^ were sacrificed 4 weeks after the beginning of the tamoxifen treatment. Regarding the high sugar tolerance tests, mice were challenged (i) either with solid diets composed of high-sucrose (60% sucrose, 20% casein, 7,95% cellulose, 7% soja, mineral, vitamin, cystin and bitartrare choline) or high-lactose (60% lactose, 20% caseine, 7,95% cellulose, 7% soja, mineral, vitamin, cystin and bitartrare choline) (SAFE^®^) during 4 or 40 days, (ii) or with 60% glucose (G8270, Sigma), 60% fructose (F9048, Sigma) diluted in water or 60% galactose diluted in water and administrated at 1% of body weight (G5388, Sigma) added in the drinking water for 3 days. Body weight was measured on a daily or weekly basis and mice were sacrificed by cervical dislocation. In order to obtain a mouse model where enteroendocrine L cells are deficient for ChREBP, the transgenic GLU**-**Venus mice, expressing Venus yellow fluorescent protein under the control of the proglucagon gene (Reimann et al. 2008), were crossed with Ch^+/+^ or Ch^−/−^ mice.

Male and female C57BL/6J mice of approximately 3 months of age, or 8-week-old *ob/ob* mice purchased from Charles River were used to generate offspring used in this study. Mice were housed on a 12-hour dark-light cycle with free access to water and standard diet. For suckling/weaning transition studies, adult male and female B6 mice were pair housed and females were monitored daily and singly housed within 48 h of parturition. 8-day-old suckling pups and 28 day-old weaned mice were then sacrificed by cervical dislocation before intestine collection. In another set of experiments, type 1 diabetes was induced by a single injection of streptozotocin (Sigma, 101809717) in 8-week-old male C57BL/6J mice. Mice were fasted 6h prior to the STZ i.p. injection and the control mice were injected with citrate buffer. The STZ was dissolved in citrate buffer (pH = 4.5) at a dose of 180mg/kg body weight. Random blood glucose (RBG) was measured before and 7 days after citrate or STZ administration.

### Indirect calorimetric measurements

Mice were housed individually in metabolic cages (Labmaster, TSE systems GmbH) with ad libitum access to food and water. They were acclimated four days before monitoring for the next three days. Oxygen consumption, carbon dioxide production (ml/h), energy expenditure (EE; kcal/h) and respiratory exchange ratio (RER; vCO2/vO2) were measured using indirect calorimetry. Subsequently, each value was expressed either by total body weight or whole lean tissue mass determined by TD-NMR (Minispec LF90II, Bruker) Food and water intakes as well as spontaneous locomotor activity were recorded during the entire experiment.

### Glucose and insulin tolerance tests

For oral and i.p. glucose tolerance test (OGTT and IPGTT, respectively), overnight fasted (16 h) mice received a 2 g/kg glucose load. For insulin tolerance test (ITT), mice fasted for 6 h were i.p. injected with 1 U/kg insulin. Blood glucose was then measured at the tail vein with an Accu-Check glucometer (Roche Diabetes Care). Intestinal glucose absorption was estimated by calculating the glycemic slope between 0 and 5 min after an oral glucose load.

### Gastric emptying measurement

Mice were fasted 24 hours with free access to water until 3 hours before an oral load with 150 µl of prewarmed (37°C) phenol red 0,5% (Sigma) in 1.5 % methylcellulose. Control mice were sacrificed immediately after the administration (100% phenol red remaining in the stomach 0 min) and the others were sacrificed 5, 30, 60 or 120 minutes after the oral load. A laparotomy was performed to expose the stomach that was ligatured at both extremities (pylorus and esophageal sphincter) before being sampled. The stomach and its content were homogenized in 30 ml NaOH (0.1N). After 1-hour decantation at room temperature, 5 ml of supernatant was added to 0.5 ml trichloroacetic acid solution (20% w/v) to precipitate the proteins. The mixture was centrifugate 10 minutes at 2500 g and the supernatant was added to 4 ml NaOH (0.5 N) to develop the maximum color intensity. The optical density (OD) was measured at 560 nm. The percentage of gastric emptying was calculated as follow: (1 – phenol red remaining in the stomach / average of phenol red in the control stomachs) * 100.

### Intestinal transit time

Carmine red was given by gavage to mice fasted for 6 h (10 mg/mL of water, 10 µL/g body weight). The total intestinal transit time was measured by determination of time between ingestion of carmine red and first appearance of the dye in feces.

### Intestinal permeability analysis in vivo

*In vivo* intestinal permeability was evaluated by the intestinal permeability of FITC-dextran 4 kDa. Briefly, 6-h water-fasted mice were gavaged with FITC-dextran 4 kDa by gavage (600 mg/kg body weight, 120 mg/ml; Sigma-Aldrich, St Louis, MO). After 4 h, 120 µl of blood were collected from each mouse from the retro-orbital vein. The blood was centrifuged at 4 °C, 10,000 rpm, for 5 min. Plasma was analyzed for FITC-dextran 4 kDa concentration with a fluorescence spectrophotometer (SPARK 10M, TECAN) at excitation and emission wavelengths of 485 nm and 535 nm, respectively. Standard curves for calculating the FITC-dextran 4 kDa concentration in the samples were obtained by diluting FITC-dextran 4 kDa in PBS.

### Imaging intestinal glucose absorption and biodistribution using 2-FDG-PET-CT

CTL and Ch^ΔGUT^ mice were fasted overnight and subjected to Positron Emission Tomography - Computed Tomography (PET-CT) using the radiotracer 2-deoxy-2-[^18^F]fluoro-D-glucose (2-FDG) to assess intestinal glucose absorption and biodistribution in peripheral tissues. Mice were anesthetized with 1–2% isoflurane (IsoVet 100%, Centravet) in 100% O_2_ during the whole experiment. PET acquisitions were acquired in the PET-CT scanner (nanoScan PET-CT, Mediso) 1 min after oral gavage of 10 MBq [^18^F]2-FDG (Gluscan, Advanced Applied Applications) in a saline solution containing 4 g/kg of D-glucose. Body temperature and respiration were registered. List-mode PET data were collected between 1- and 60-min post-injection, binned using a 5-ns time window, a 400- to 600-keV energy window, and a 1:5 coincidence mode. At the end of the PET acquisitions, CT scans were performed using the following parameters: mode semi-circular, tension of 39 kV, 720 projection full scan, 300 ms per projection, binning 1:4. After the *in vivo* scans, mice were sacrificed and the whole intestine was isolated and flushed to remove intraluminal 2-FDG for *ex vivo* PET-CT acquisition of 5 min. *In vivo* PET acquisitions were reconstructed in 12 frames of 5 min using the Tera-Tomo reconstruction engine (3D-OSEM based manufactured customized algorithm) with expectation maximization iterations, scatter, and attenuation correction. Images were analyzed using the software PMOD (PMOD Technologies LLC). Standardized Volume of Interest (VOI) was drawn in each organ, and Standardized Uptake Values (SUV) were calculated by dividing the mean tissue radioactivity concentration by the total orally administrated 2-FDG and body weight.

### Plasma β-hydroxybutyrate and insulin measurement

Plasma β-hydroxybutyrate concentrations were measured at the tail vein of fasted and refed mice with a ketone meter (FreeStyle, Optium Neo). Plasma insulin was assayed using an ultrasensitive insulin ELISA kit (ALPCO). To that end, blood samples were collected by retro-orbital sinus puncture at 0, 15 and 30 min of OGTT and immediately centrifuged (3000g, 4°C, 10min) before storage at −80 ° C for further analysis.

### Measurement of portal and gut mucosa GLP-1 or GIP levels

Portal GLP-1 levels were measured before and 15-min after an oral challenge with 2 g/kg glucose in overnight fasted mice. Blood was collected from the portal vein of anesthetized mice (ketamine 100 mg/kg, xylazine 10 mg/kg i.p.) in EDTA-precoated tubes containing DPP-IV inhibitor (DPP4-010, 5mM, Millipore^TM^), aprotinin (5mM, Sigma®) and protease inhibitor cocktail (P8340, Sigma®). Blood samples were then immediately centrifuged (3000g, 4°C, 10min) and stored at −80°C. For GLP-1 and GIP gut content evaluation, jejunum from 16 hours fasted mice was collected and sliced into small pieces. Tissues were homogenized in ethanol/acid (100% ethanol: sterile water: 12N HCl 74:25:1 v/v) solution (5 mL/g tissue) and homogenates were shred (6,5 movements/sec, 60 sec) and incubated overnight at 4°C. The next day, samples were centrifuged (3000g, 20 min, 4°C) and supernatants were collected and stored at −80°C until GLP-1 dosage. Portal and gut mucosa GLP-1 or GIP levels were assayed using high sensitivity GLP-1 total chemiluminescent ELISA kit (K150JVC-1, Meso Scale Discovery®) or GIP total ELISA kit (EZRMGIP-55K, Sigma-Aldrich, Merck).

### Ex vivo intestinal glucose absorption

Glucose transport was assayed *ex vivo* using jejunal loop as described previously.1 Briefly, four 3-cm intestinal segments were filled with Krebs Ringer Bicarbonate Buffer solution containing 30 mM D-glucose with 0.1 mCi/mL [^14^C]-glucose (specific activity 49.5 mCi/mmoL) and with or without 100 mM phloretin, a glucose transporter inhibitor. Each segment was ligated at both ends and incubated in a 37°C thermostat-controlled bath of Krebs modified buffer at pH 7.4 continuously gassed with 95% O25% CO2. Mucosalto-serosal and serosal-to-mucosal transport of glucose was monitored using everted and noneverted isolated intestinal loops, respectively. Time-dependent [14C]-glucose transport was determined by sampling from the bath at 0, 5, 10, 20, 30, and 60 minutes. At 60 minutes, isolated intestinal loops were collected, flushed, weighed, and homogenized with Ultra-Turrax (Ika, Wilmington, NC) for quantification of radioactivity. Radioactivity was measured using a beta counter (Beckman LS 6000 TA liquid scintillation counter). Apparent permeability (Papp) was used to assess transport according to the following equation Papp ¼ (dQ / dt) $ (V / Q0 $ A), where V is the volume of the incubation medium, A is the area of the loop, Q0 is the total radiolabeled glucose introduced into the loop and dQ/dt is the flux across the intestinal loop.

### Isolation of intestinal epithelial cells (IEC)

IEC from 8-week-old C57BL/6J were isolated by incubation of small intestinal or colonic fragments in a PBS chelating buffer containing EDTA (15 mM), Dithiothreitol (DTT, 1M) and protease inhibitors (1 tablet per 50mL extraction solution) (COmplete protease inhibitor cocktail tablets in EASYpacks, Roche®) for 30 min at 37°C under gentle shaking at 100 rpm. Cells were pelleted (8000g, 4°C, 5 min) and flash-frozen in liquid nitrogen and stored at −80 °C until further protein and gene expression analyses. The sequential isolation of intestinal and colonic epithelial cells along the crypt–villus axis (CVA) was performed as previously described and validated (Guilmeau et al., 2010). The small and large intestines were dissected, everted, filled to distension with phosphate-buffered saline (PBS), and incubated with shaking at 37°C in 1.5 mM EDTA buffer. Resulting fractions of dissociated epithelial cells were harvested by centrifugation at 1500 rpm at 4°C for 5 minutes, cell pellets were snap frozen in liquid nitrogen, and stored at −80°C.

### L cells isolation and flow cytometry analyses

IEC isolation was carried out from 10- to 12-week-old male GLU-Venus mice. After cervical dislocation, proximal jejunum and colon were collected, washed in PBS and mesenteric adipose tissue and muscular layers were separated from the epithelium. Epithelial tissue were opened longitudinally, cut into 1-2 mm pieces and digested 30 min at 37°C in DMEM containing 25 mM of glucose (G25) and 1M of collagenase (Clostridium, C9407-1G, Sigma®) under repeated agitation every 10 min. Sedimented cells at the bottom of the tube were placed in a new collagenase solution for a second digestion step while the supernatant was centrifuged (1600 rpm, 10 min). After 30 min epithelial cells suspension were collected by centrifugation (1600 rpm, 10 min), pooled in DMEM (G25) and passed through large (70/100 µm) and then small diameter (40 µm) filters. All isolated epithelial cells were pelleted through centrifugation (1600 rpm, 5 min) and resuspended in DMEM (G25) before Fluorescence-activated cell sorting (FACS, 488 nm excitation) using an ARIA III high flow sorter analyzer (Beckman Coulter®). Single cells were selected by lateral dispersion, direct scattering and pulse width in order to exclude cellular debris and aggregates. L^+^ cells expressing the Venus protein (L^+^) were discriminated from L^-^ cells by their relative fluorescence, allowing the collection of isolated L^+^ cells at ∼ 95% purity. The mixed population of other epithelial L^-^ cells was also collected apart. Up to 2 000 for L^+^ and 50 000 for L^-^ cells were sorted into 200 µL of TRIZol and frozen at −80°C for subsequent RNA extractions.

### GLUTag cell culture and transfections

The GLUTag mouse enteroendocrine cell line was kindly provided by Daniel J Drucker (Mount Sinai Hospital, Toronto, ON, Canada). GLUTag cells were grown at 37°C under 5% CO_2_ at 95% humidity in Modified from Dulbecco Medium (DMEM) plus GlutaMAX^TM^-1 medium containing glucose (5,5mM) with glutamine and sodium pyruvate (Thermo Fisher Scientific), supplemented with fetal bovine serum (10%, FBS), penicillin (1%, 10 000U/ml) and streptomycin (10 000 µg/ml). For glucose challenges experiments, GLUTag cells were starved overnight in a medium containing low glucose (1mM), glutamine but no sodium pyruvate (Thermo Fisher Scientific), supplemented with FBS (1%), and incubated for 24h with either low (5,5mM) or high glucose (25mM), supplemented or not with SBI-477 (10µM). Supernatants were then collected, centrifuged (5 min, 1000 rpm), transferred to fresh Eppendorf tubes and stored at-80°C for further analysis.

GLUTag cells were plated in 24-well dishes, allowed to recover for 48h before transfection. siRNAs and plasmids transfections were performed using OptiMEM media and Lipofectamine 2000 as instructed by the manufacturer (Invitrogen). The cells were incubated with the transfection mixture for 4 h, washed twice with DMEM containing 5.5 mM glucose supplemented with FBS (1%) and then cultured in regular growth medium for 24, 48 or 72  h, and finally harvested and stored at −80  °C. Either non specific siRNA (ns) or siRNA targeting murine ChREBP (siChREBP) (SMARTPool, ON-TARGETplus Mouse Mlxipl siRNA, Dharmacon) were used. *Gcg*-Luc luciferase reporter plasmids, in which luciferase activity was placed under the control of −2400 bp of the rat *Gcg* gene promoter, were provided by Dr Tianru Jin (University of Toronto, Toronto, Canada) (Lü et al., 1996). ChoRE-luc reporter, in which luciferase activity was placed under multimeric ChoRE sequences of the rat *L-pk* gene promoter, was a gift from Dr Mireille Vasseur (Bricambert et al., 2018). The Renilla luciferase reporter pRL-CMV was used as internal control. Dual luciferase reporter assays were performed 48h post transfection. Empty pcDNA3 plasmid, ChREBP-WT and ChREBP-CA, encoding mouse wild type ChREBP or a constitutively active form of ChREBP deleted of the LID domain were previously described (Iroz et al., 2017).

### Protein extraction and western blot analyses

IECs from small intestine and colon were solubilized at 4°C in a buffer containing NaCl (150mM), Tris-HCl 1M (50mM, pH 7,5), EDTA 0,5M (5mM), Sodium Pyrophosphate (30mM), NaF (5mM), triton X-100 (1%), PMSF (1mM), Sodium Orthovanadate (2mM) and protease inhibitors tablets (COmplete protease inhibitor cocktail tablets in EASYpacks, Roche). Supernatants were collected after centrifugation (14000g, 4°C, 10 min) and proteins were quantified using the Bradford method (BioRad Protein Assay). Whole-cell lysates (80 microgram) were subjected to SDS-PAGE electrophoresis and immunoblotted with the following primary antibodies: anti-ChREBP (1:1000, #NB-135, Novus Biologicals), anti-PCNA (1: 1000, #13110, Cell Signaling Technology) and anti-β-actin (1:5000, #13E5, Cell Signaling Technology). The immunoreactive bands were revealed using Clarity Western ECL Substrate (BIO-RAD). Chemiluminescence analyses were performed with the ChemiDoc MP Imaging System (BIO-RAD).

### Histological analyses and GLP-1 immunostaining

The small intestine and the colon were collected, opened longitudinally, cleaned with PBS (Sigma®), rolled up and then fixed in paraformaldehyde solution (4%, PFA) all night at 4°C. The tissues were embedded in paraffin and cut into 5 µm thick sections before deparaffinization and rehydration. Transversal intestinal sections from CTL and Ch^ΔGUT^ mice were stained with hematoxylin and eosin to visualize crypt and villus morphology. On intestinal sections from Ch^+/+^ and Ch^-*/*-^ mice, endogenous peroxidase activity was blocked by H_2_O_2_ (3%, 10 min) (H-1009, Sigma®) and L-cells immunolabelling was performed using anti-GLP-1 antibody (1:200, ab26278, Abcam®), mouse on mouse immuno-detection kit (BMK-2202, M.O.M^TM^, Vector®), ABC HRP kit (PK-6100, The VECTASTAIN® Elite, Vector®) and DAB Peroxidase HRP Substrate kit (SK-4100, Vector®) according to manufacturer’s instructions. Following dehydration and counterstaining with hematoxylin, sections were mounted in VectaMount Permanent Mounting Medium (H-5000, Vector®), scanned with the Lamina Multilabel Slide scanner (Perkin Elmer) and analyzed with the “Panoramic Viewer” software. Total GLP-1 positive L cell number was evaluated manually in a blinded way on a single transversal section of intestinal roll from the upper small intestine (USI), the lower small intestine (LSI) and the colon (COL).

### Transmission electron microscopy

Samples from jejunum were analyzed under a JEOL 1011 (Tungsten filaments) transmission electron microscope with a charge-coupled device (CCD) digital camera mounted in the 35mm port on the microscope (GATAN Erlangshen). Acquisitions were processed using software « Digital Micrograph » and ImageJ. The quantitative analysis (microvillus length and density) was obtained from 10 cells of each sample (40 cells by group).

### Quantitative RT-PCR analyses

Total RNA was extracted from mouse small intestine and/or colon using TRIzol reagent (15596026, Invitrogen^TM^) according to the manufacturer’s instructions and RNA purity was verified by spectrophotometry (Nanodrop^TM^ 3300, Thermo Scientific®). cDNA was obtained by retro-transcription of 1µg of purified RNA using Super Script III Reverse Transcriptase kit (Invitrogen^TM^). Quantitative PCR was carried out in duplicate using SYBR Green I Master (Roche®) and LightCycler480 System (Roche®) under following conditions: 15-min denaturation at 95°C, followed by 10 s cycles at 95°C (denaturation), 45 s at 60°C (annealing), and 10 s at 72°C (extension). Primers sequences used are detailed in Table 1. Gene expression was normalized over the expression of *TBP* mRNA levels.

**Table 1:**
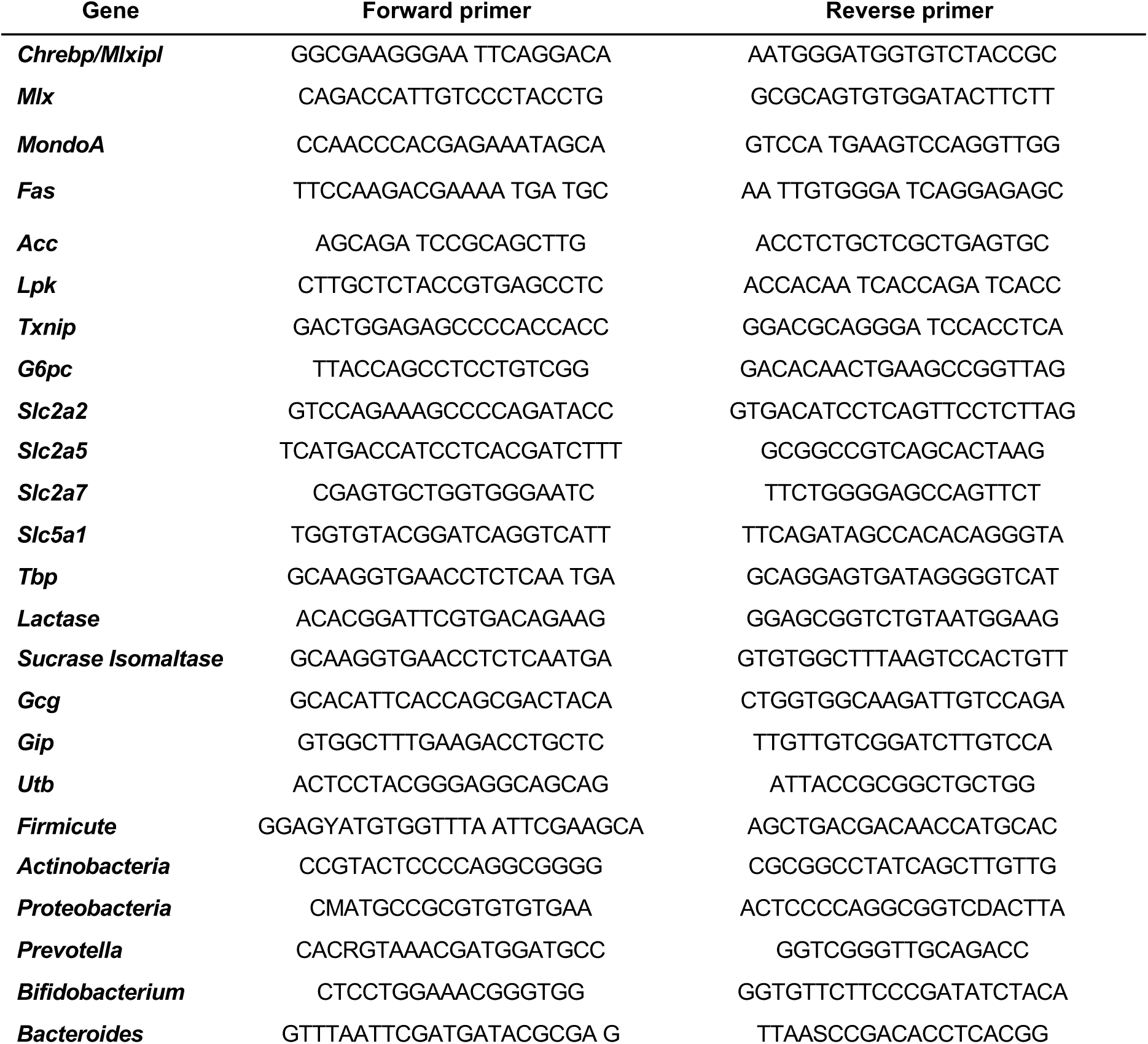
Primers sequences used for PCR analysis

### Cecal microbiota analysis

Bacterial DNA was extracted from luminal cecal content samples using the ZR fecal DNA Miniprep kit according to the manufacturer instructions (Zymo Research, USA). Relative abundance of bacterial phyla or species were determined by PCR analysis using the primers indicated in Table 1 and each value was normalized to *UTB* levels. Quantitative PCR was carried out in duplicate using SYBR Green I Master (Roche®) and LightCycler480 System (Roche®) under following conditions: 15-min denaturation at 95°C, followed by 10 s cycles at 95°C (denaturation), 45 s at 60°C (annealing), and 10 s at 72°C (extension).

### Transcriptomic analysis

Gene expression profile of jejunum epithelial cells isolated from Ch^ΔGUT^ and CTL mice in response to glucose/fructose challenge were analyzed using Affymetrix Clariom S Mouse microarrays by the “Genomics” platform of the Cochin Institute. Differential expression was measured with moderated t-test (limma R package). The gene set enrichment analyses (GSEA) were performed with these data and with BioCarta and Kyoto Encyclopedia of Genes and Genomes (KEGG) gene sets. Genes with a fold change (converted into log_2_) ≥ 1,3 or ≤ 1,3 and *P value* (converted into log10) < 0,01 were considered as differentially expressed between CTL *vs* Ch^ΔGUT^ mice in response to 20%-glucose challenge and were analyzed by Partek Genomics Suite (Version 7-19-1125), to generate volcano plot. Differential gene expressions levels with a fold change ≥ 1,3 or ≤ 1,3 and *P value* < 0,01 between CTL untreated (water without sugar) *vs* CTL treated (20%-glucose challenge mice) and CTL *vs* Ch^ΔGUT^ mice both in response to 20%-glucose challenge were analyzed using the IPA software (current version 51963813, release 2020-03-11, Ingenuity Systems Inc., QIAGEN, Redwood City, CA, USA) by uploading outcomes from microarray to identify statistically overrepresentation of Gene Ontology (GO) terms in gene sets. VENNY^2.1^ was used to generate Venn diagram by uploading outcomes from RNA sequencing. FunRich (Functional Enrichment analysis tool) software has also been used to create complex venn diagram. Genes were considered as differentially expressed when fold change was ≤ 1,3 were analyzed by Heatmapper, a web server freely available, to generate a selected heatmap from transcriptomic analysis.

### Statistical Analysis

All values are presented as means ± SEM. Data were analyzed using one-way or two-way analysis of variance (ANOVA) followed by Bonferroni post-test multiple comparisons or Mann-Whitney U test. Data analysis was performed using GraphPad Prism software. Statistical significance was assumed at *P* values <0.05.

## RESULTS

### Tolerance to oral glucose challenge is paradoxically improved in Ch^−/−^ mice

We first evaluated the metabolic consequence of ChREBP deficiency. We observed that despite higher fasting blood glucose, insulin resistance and intolerance to peripheral glucose compared to Ch*^+/+^* control mice (Figure 1A-D), Ch*^−/−^* mice surprisingly displayed an improved tolerance to oral glucose challenge (Figure 1E-F). In agreement with previous publications (Iizuka et al., 2004), ChREBP deficiency was accompanied by a 25% reduction in white adipose tissue weight and a 75% increase in liver mass despite similar body weight and length in Ch*^−/−^* and Ch*^+/+^* mice (Figure S2A). We next monitored Ch^-/-^ mice in metabolic cages for 5 consecutive days. No significant change in food, water intake, respiratory exchange ratio (RER), oxygen consumption rate (VO2), carbon dioxide release rate (VCO2) or body heat production (Figure S2B-G) could be observed. Of note, the locomotor activity of Ch*^−/−^* mice was significantly reduced during the nocturnal period when compared to control littermates (Figure S2I-J). We validated that the paradoxical ameliorated response of Ch*^−/−^* mice to OGTT did not result from slower gastric emptying upon ChREBP deficiency (Figure S2J). Tissue distribution analysis revealed abundant *mlxipl* gene expression in the upper small intestine, where carbohydrates digestion and absorption primarily occur (Figure 1G). Indeed, quantification of *mlxipl* mRNA demonstrated that duodenal expression represented 57% of that in the liver, a major ChREBP expressing tissue (Iizuka et al., 2004). Gut *mlxipl* mRNA levels followed a decreasing gradient along the antero-posterior axis, illustrated by the 9.6 fold drop in the expression of *mlxipl* when comparing the expression in the duodenum and the colon. A smaller range gradient and (3.5 fold difference in duodenum *vs* colon) and an opposite distribution was observed for *mlx,* the obligatory partner of ChREBP (Stoeckman et al., 2004) (Figure 1G). In agreement with mRNA quantifications, western blot analysis revealed that ChREBP protein level was predominant in the duodenum and the jejunum, and less abundant in caecum and colon (Figure 1H). Moreover, epithelial cell fractionation along the crypt-villus axis demonstrated that, despite higher *mlxipl* mRNA levels in jejunal crypts, ChREBP is both expressed in villi differentiated cells and PCNA positive proliferating cells from the crypt compartment (Figure 1J-K). Interestingly, jejunal loops exposure to luminal 25mM glucose resulted in a significant increase of epithelial intestinal *mlxipl* and *mlx* mRNA levels, as well as ChREBP protein levels as compared to those stimulated with 5mM glucose (Figure 1L-N). In parallel, exposure of jejunal loops to 25 mM glucose also led to enhanced expression of prototypic ChREBP target genes such as *txnip* and *lpk* suggesting that the ChREBP signaling pathway was fully functional in the gut epithelium (Figure 1O). Of note, intestinal mRNA levels of lipogenic genes (*acc, fas, scd-1*) that are known to be controlled by ChREBP in the liver, were not modified in response to either high glucose challenge or ChREBP deficiency (Figure S2K). Finally, we verified that loss of ChREBP in the gut epithelium was not compensated by increased expression of *mondoA*, its paralog protein (Figure S2L). Altogether, these data suggest that improved tolerance of Ch*^−/−^* mice to oral glucose bolus may result from impaired ChREBP activity in the gut mucosa.

**Figure 1:**
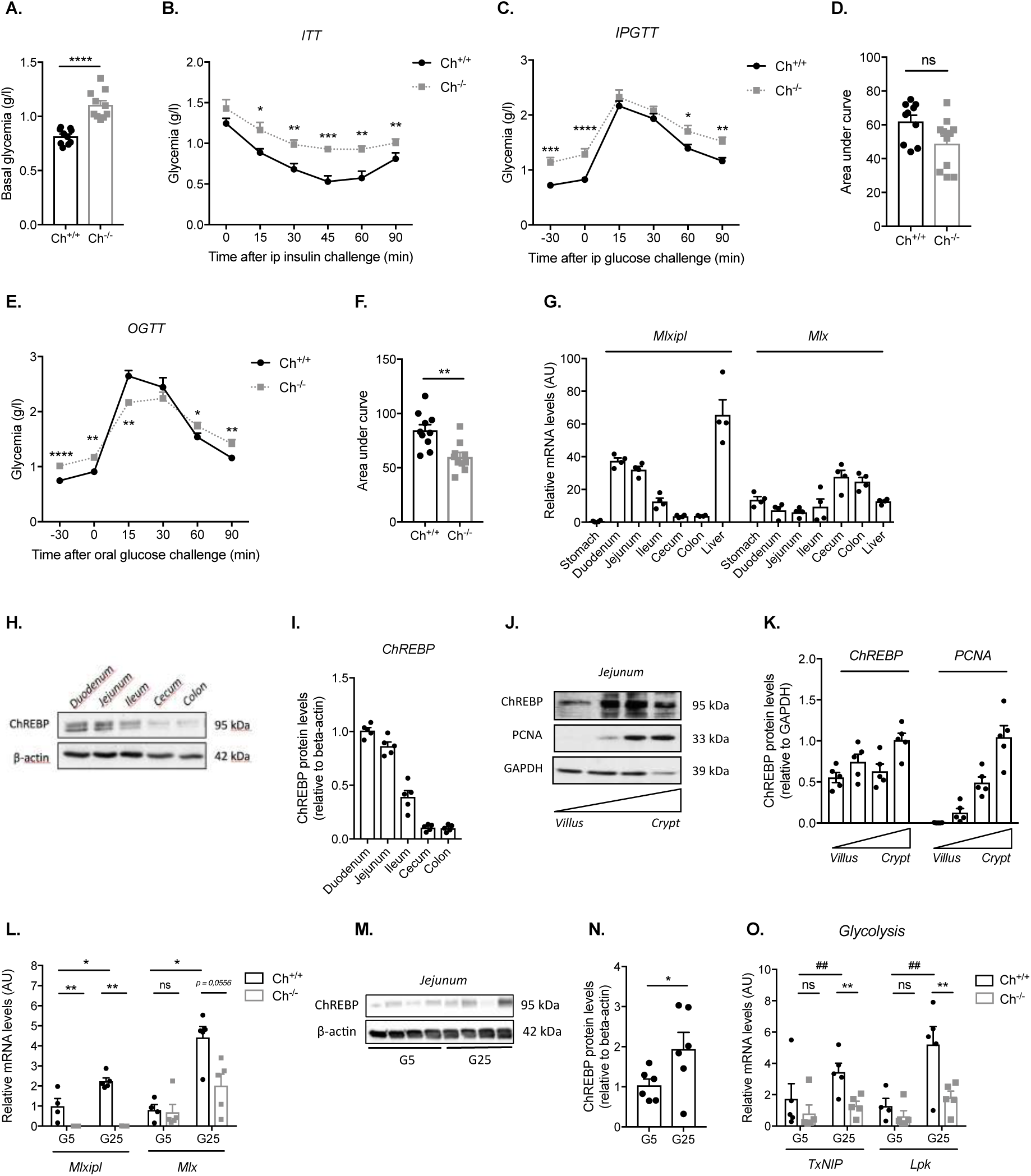
Whole body ChREBP deficiency improves oral glucose tolerance. **(A)** Plasma glucose was measured in overnight-fasted 10- to 12-week-old male wild type (Ch^+/+^) and ChREBP-KO (Ch^-*/*-^) mice (n = 10). **(B)** Insulin tolerance test (insulin i.p. 1U/kg) was performed in 10- to 12-week-old male Ch^+/+^ and Ch^-*/*-^ mice after 6h of fasting (n = 11-12). **(C-F)** Glucose tolerance test was performed following i.p. injection **(C-D)** or an oral bolus of glucose (2g/kg) **(E-F)** in overnight fasted 10- to 12- week-old male Ch^+/+^ and Ch^-*/*-^ mice and **(D, F)** corresponding area under curves (AUC) was calculated (n = 10). **(G)** Relative mRNA levels *mlxipl* and *mlx* (normalized against *TBP)* in isolated epithelial cells from the stomach and the gut and in the liver of 10- to 12-week-old male wild type mice (n = 4). **(H-K)** Representative western blot analyses **(H, J)** and corresponding densitometric quantifications **(I, K)** of ChREBP protein levels in epithelial cells isolated from various intestinal segments **(H-I)** or from the jejunum along the crypto-villus axis **(J-K)** of 10- to 12-week-old male wild type mice (n = 5). β-actin was used as a loading control and PCNA as a validation of cell fractionation. **(A-K)** Values are means ± SEM. *\*P < 0.05, **P < 0.01, ***P < 0.001, ****P < 0.0001*; compared Ch^+/+^ *vs* Ch^-*/*-^ mice (Mann Whitney U test). **(L-O)** Epithelial cells isolated from jejunal loops of overnight fasted Ch^+/+^ and Ch^-*/*-^ mice treated *ex vivo* for 4h with 5mM or 25mM glucose. **(L, O)** Relative mRNA levels of *mlxipl, mlx, txnip, and lpk* (normalized to *TBP*) and **(M)** ChREBP western blot analyses and **(N)** corresponding protein levels quantifications of ChREBP protein levels (n = 5). β-actin was used as a loading control. **(L-O)** Values are means ± SEM. *^##^P < 0.01;* compared G5 *vs* G25 and *\*\*P < 0.01*; compared Ch^+/+^ *vs* Ch^-*/*-^ mice (Mann Whitney U test).

### Gut epithelial ChREBP activity contributes to glycemic control

In order to determine whether intestinal ChREBP activity contributes to the improved oral glucose tolerance observed in Ch^-/-^ mice, we generated a mouse model of inducible deletion of ChREBP in the gut epithelium (Ch^ΔGUT^ mice). Efficient intestinal ChREBP deficiency was validated 4 weeks after tamoxifen administration in 8-week-old mice, as illustrated by the drastic reduction in *mlxipl* mRNA levels in epithelial cells isolated from the upper small intestine (USI), the lower small intestine (LSI) and the colon of Ch^ΔGUT^ mice as compared to CTL mice (Figure 2A). Accordingly, ChREBP protein levels were undetectable in intestinal and colonic epithelial cells of Ch^ΔGUT^ mice, while ChREBP expression was unchanged in the liver, validating the specificity of the invalidation in the gut epithelium (Figure 2B). Of note, despite normal body weight, fasted normoglycemia, unchanged tolerance to intraperitoneal glucose challenge and unmodified insulin sensitivity, Ch^ΔGUT^ mice displayed improved tolerance to an oral glucose load compared to CTL mice (Figure S3A, 2C-E). A significant and early reduction of glycemic excursion was observed in Ch^ΔGUT^ mice at fifteen minutes of OGTT (Figure 2F). Examination of the gut demonstrated no difference in weight or length of the small intestine or the colon in Ch^ΔGUT^ mice compared to CTL littermates, suggesting that loss of local ChREBP activity had no impact on gut epithelium trophicity (Figure S3B-E). Moreover, specific inactivation of ChREBP in the gut did not alter intestinal or colonic epithelial histology, as illustrated by proper crypt-villus organization as well as conserved crypt and villus length (Figure 2G). However, functional analysis revealed that bowel transit time of Ch^ΔGUT^ mice was faster (a 13.2% increase in time) and that their relative gut epithelial permeability was reduced (a 39.5% reduction), compared to CTL mice (Figure 2H-I). Altogether, these results highlight that loss of gut epithelial ChREBP activity contributes to glycemic control by reducing blood glucose raise upon glucose luminal challenge specifically.

**Figure 2:**
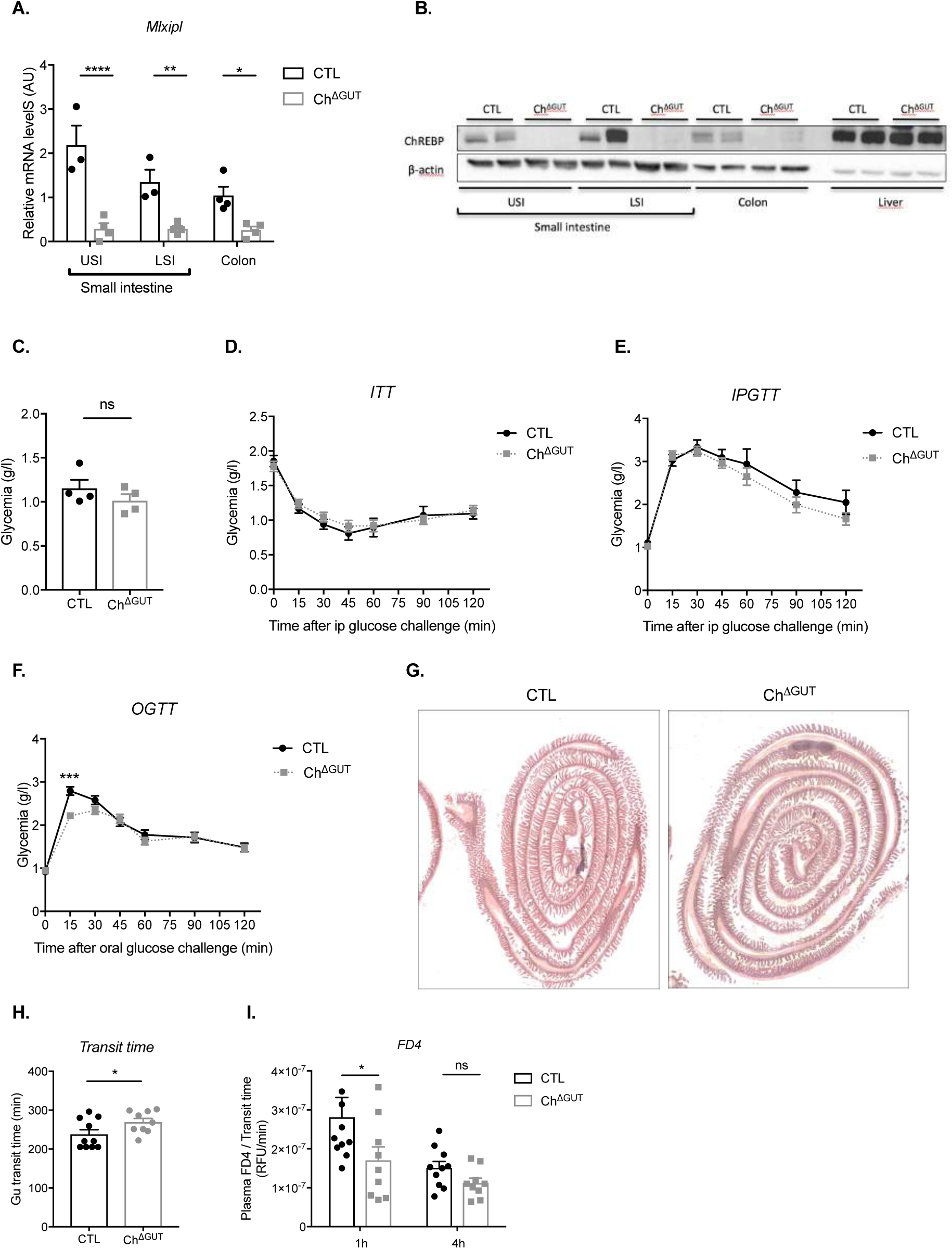
Intestinal ChREBP activity contributes to glycemic excursion upon an oral glucose load. **(A)** Relative *mlxipl* mRNA levels in intestinal and colonic epithelial cells isolated from 16-week-old male control (CTL) or gut-specific ChREBP-KO (Ch^ΔGUT^) mice (n = 3-4). Intestinal segments from Upper Small Intestine (USI), Lower Small Intestine (LSI) and Colon (COL) were treated *ex vivo* for 4h with 25mM glucose before epithelial cell isolation. Values were normalized against *TBP* and represent means ± SEM. *\*\*P < 0.01*; compared CTL *vs* Ch^ΔGUT^ mice (two-way ANOVA followed by Bonferroni correction for multiple comparisons). **(B)** Representative western blot analysis of ChREBP protein levels on epithelial cells isolated along the gut and in the liver of 16- week-old male CTL and Ch^ΔGUT^ mice. β-actin was used as a loading control. **(C)** Plasma glucose of overnight-fasted 16-week-old male CTL and Ch^ΔGUT^ mice (n = 4). **(D)** Insulin tolerance test (i.p. insulin 1U/kg) was performed in 16-week-old male CTL and Ch^ΔGUT^ mice after 6h of fasting (n = 15-21). **(E-F)** Glucose tolerance tests were performed in 16-week-old male CTL and Ch^ΔGUT^ mice after 16 h of fasting followed by an i.p. injection **(E)** or an oral bolus **(F)** of glucose (2g/kg) (n = 16-23). **(C-F)** Values are means ± SEM. *\*\*\*P < 0.001*, *\*P < 0.05, *P < 0.05;* compared CTL *vs* Ch^ΔGUT^ mice (Mann Whitney U test). **(G)** Hematoxylin & eosin staining of small intestinal transversal sections of 12-week-old male CTL and Ch^ΔGUT^ mice**. (H)** Total intestinal transit was evaluated by the time required for expelling the first red pellet following oral gavage with carmine red dye (n = 9-10). **(I)** *In vivo* intestinal permeability in 16-week-old male CTL and Ch^ΔGUT^ mice, as quantified by plasma FD4 at 1h and 4h after the oral administration of the tracer (n = 9-10). Data are reported as relative fluorescent units (RFU). **(H-I)** Values are means ± SEM. *\*\*\*P < 0.001*, *\*P < 0.05, *P < 0.05;* compared CTL *vs* Ch^ΔGUT^ mice (Mann Whitney U test).

### The loss of intestinal ChREBP impairs GLP-1 production

We next addressed whether the improved OGTT upon gut ChREBP deficiency (Figure 2D) could be due to a modification of the incretin response. While intestinal *gip* mRNA levels and mucosal GIP content were not significantly changed upon gut ChREBP deficiency (Figure 3A-B), we evidenced that production of GLP-1, another key incretin hormone, was impaired in Ch^ΔGUT^ mice as compared to CTL mice (Figure 3C, E-F). Indeed, despite similar fasting portal GLP-1 concentrations both groups of mice, the stimulatory effect of an oral glucose challenge (15 min time point) on GLP-1 secretion was blunted in Ch^ΔGUT^ mice (Figure 3C). Of note, plasma insulin concentrations remained however unchanged in Ch^ΔGUT^ mice as compared to CTL mice (Figure 3D). This altered GLP-1 secretory response to glucose was associated with a marked reduction in total GLP-1 content and a 62.7% decrease in *gcg* mRNA levels in the small intestine (Figure 3E-F), suggesting that ChREBP deletion dampens glucose stimulated GLP-1 secretion, at least partly through the inhibition of *gcg* transcription. These results were also confirmed in Ch*^−/−^* mice. Similarly, we observed that Ch*^−/−^* mice exhibit decreased portal GLP-1 concentrations and intestinal total GLP-1 content at 15 min after an oral glucose challenge compared to CTL mice (Figure S4A-B) as well as reduced intestinal *gcg* mRNA levels and L cells number (Figure S4C-E). We next performed GLP-1 producing L cells sorting from GLU-Venus reporter mice, which express the Venus fluorescent protein under the control of the *gcg* gene (Reimann et al., 2008) (Figure 3G). As validated by the specific expression of *gcg* and *pax6* in Venus positive L cells (Figure 3H, S4F), we demonstrated that *mlxipl* expression was enriched by 12 and 5 fold in intestinal L^+^ cells and colonic LC^+^ cells respectively, as compared to L^-^ and LC^-^ cells (Figure 3I). Of note, no significant change in *mlx* and *mondoA* mRNA levels was detected in L cells when compared to other Venus negative epithelial cells (Figure S4G-H). Significant downregulation of *gcg* mRNA levels specifically in jejunal L^+^ cells from Ch*^−/−^* mice relative to L^+^ cells from Ch*^+/+^* mice, suggested that blunted *gcg* mRNA levels in Ch^ΔGUT^ mice resulted, besides reduced L cell number, from decreased transcriptional activity of the *gcg* promoter in intestinal but not in colonic L cells (Figure 3J). To confirm that ChREBP is a direct regulator of *gcg* expression, we conducted a series of experiments *in vitro* using the GLP-1 producing GLUTag cell line. Pharmacological inhibition of ChREBP activity by SBI477 treatment (Ahn et al., 2016) in GLUTag cells prevented the glucose-mediated induction of *gcg* expression (Figure 3K). This effect was associated to reduced *mlxipl* expression (Figure S4I), and no change in *mlx* and *mondoA* mRNA levels were observed upon either high glucose concentrations or ChREBP blockade (Figure S4J-K). In addition, ChREBP silencing by siRNA approach in GLUTag cells (Figure 3L) led to 2.9 fold reduction of ChoRE-luc reporter activity (Figure 3N), and was associated with a 51.3% reduction of *gcg* mRNA levels upon high glucose concentrations (25mM) (Figure 3M). Finally, the *gcg*-luciferase reporter activity was significantly repressed upon ChREBP knock down in GLUTag cells at high glucose concentrations (25mM), pinpointing that ChREBP activity contributes to the glucose induced *gcg* transcription (Figure 3N). Conversely, overexpression of a constitutively active form of ChREBP (ChREBP CA) led to a significant increase of *gcg* promoter activity (Figure S4L). Altogether these data indicate that the incretin response is impaired in a ChREBP deficiency context and suggest that other ChREBP intestinal function contributes to the improved oral glucose tolerance in Ch^ΔGUT^ mice.

**Figure 3:**
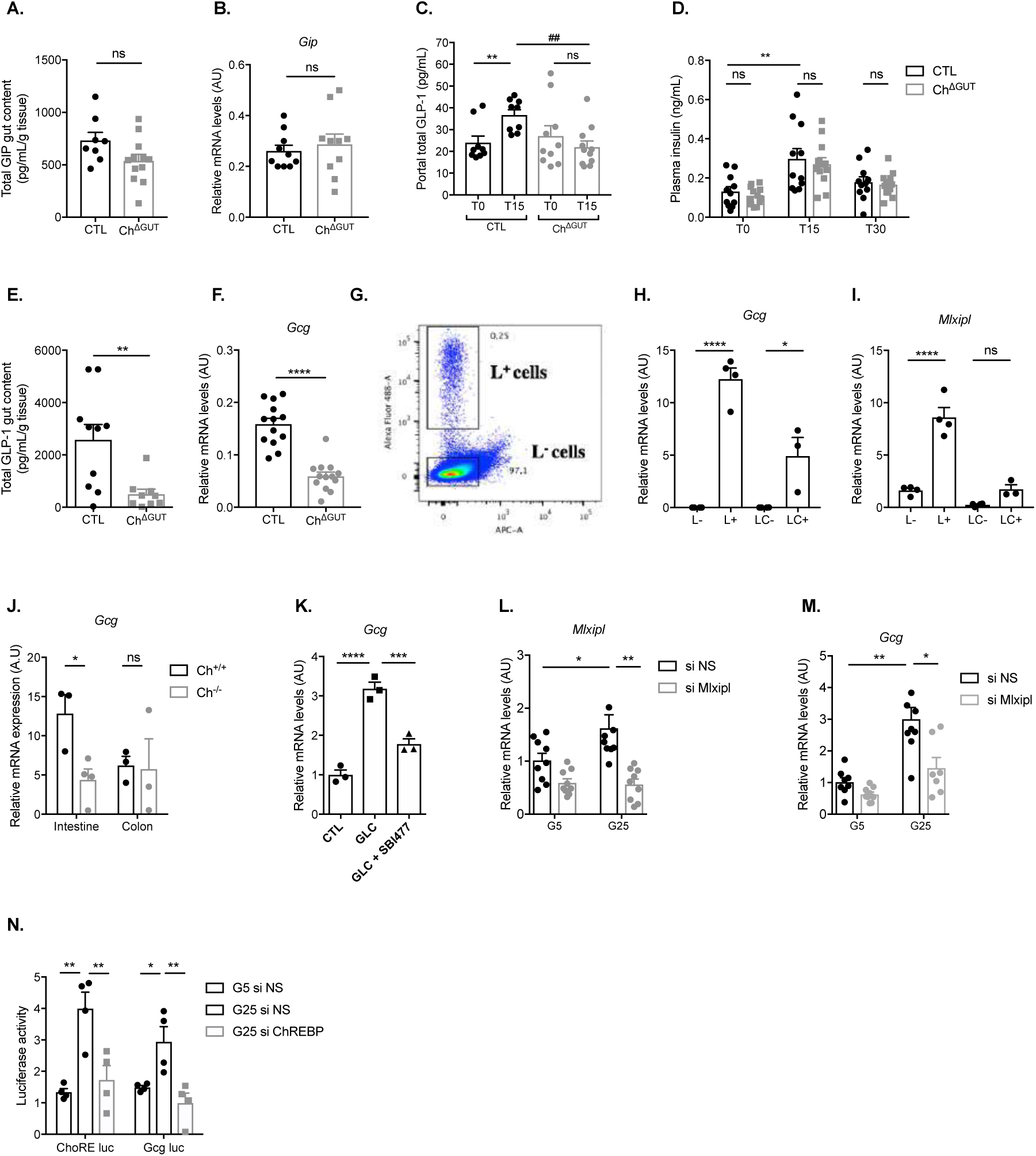
GLP-1 production is reduced upon intestinal ChREBP deficiency. **(A)** Total GIP contents measured in upper small intestinal mucosa of 16-week-old female control (CTL) and Ch^ΔGUT^ mice (n = 8-12). **(B)** Relative mRNA levels of *gip* in gut epithelial cells isolated from 16-week-old female CTL and Ch^ΔGUT^ mice (n = 10). Upper small intestinal segments (USI) were treated *ex vivo* for 4h with 25mM glucose before epithelial cell isolation. **(A-B)** Values are means ± SEM. *ns*, *\*\*\*\*P < 0.0001, **P < 0.01*; compared CTL *vs* Ch^ΔGUT^ mice (Mann Whitney U test). **(C)** Total GLP-1 concentrations measured in portal blood at 0 and 15 min after oral glucose gavage (2g/kg) of 16-week-old female CTL and Ch^ΔGUT^ mice (n = 9-11). Values are means ± SEM. *^##^ P < 0.01*; compared CTL *vs* Ch^ΔGUT^ mice; *\*\*P < 0.01;* compared T0 *vs* T15 for each genotype (two-way ANOVA followed by Bonferroni correction for multiple comparisons). **(D)** Plasma insulin concentrations at 0, 15 and 30 min after an oral challenge of glucose (2g/kg) in 16-week-old female CTL and Ch^ΔGUT^ mice (n = 11-12). **(E)** Total GLP-1 contents were measured in upper small intestinal samples at 15 min after oral glucose gavage (2g/kg) of 16-week-old female control (CTL) (black bars) and Ch^ΔGUT^ (gray bars) mice 15 min after oral glucose gavage (n = 9-10). **(F)** Relative mRNA levels of *gcg* (normalized to *TBP*) in gut epithelial cells isolated from 16-week-old female CTL and Ch^ΔGUT^ mice. Upper small intestine segments (USI) were treated *ex vivo* for 4h with 25mM glucose before epithelial cell isolation (n = 13). **(D-F)** Values are means ± SEM. *ns*, *\*\*\*\*P < 0.0001, **P < 0.01*; compared CTL *vs* Ch^ΔGUT^ mice (Mann Whitney U test). **(G)** Representative FACS analysis of jejunal cells isolated from 10- to 12- week-old male GLU-Venus mice. L+ indicates cells that were positive in the green channel (488 nm excitation laser) detecting Venus. The gated region named L-outlines Venus-negative cells. **(H-I)** Relative mRNA levels of *gcg and mlxipl* (normalized to *TBP*) in GLP-1 producing epithelial cells (jejunal L+ and colonic LC+ cells) and in GLP-1 negative epithelial cells (jejunal L- and colonic LC-), of 10- to 12- week-old male GLU-Venus mice (n = 3-4). Values were and represent means ± SEM. *ns*, *\*P < 0.05, ****P < 0.0001*; compared L^-^ *vs* L^+^ and LC^-^ *vs* LC^+^ (one-way ANOVA). **(J)** Relative *gcg* mRNA levels (normalized to *TBP*) in intestine and colon of 10- to 12-week-old male GLU-Venus and GLU-Venus Ch^-*/*-^ mice (n = 3-4). Values are means ± SEM. *ns*, *\*P < 0.05*; compared GLU-Venus *vs* GLU-Venus Ch^-/-^ mice (two-way ANOVA followed by Bonferroni correction for multiple comparisons). **(K)** Relative *gcg* mRNA levels (normalized to *TBP*) in GLUTag cells incubated 24h with 5 (G5) or 25mM glucose (G25) +/-SBI477 (10µM) (n = 3). Values are means ± SEM. *\*\*\*P < 0.001*, *\*\*\*\*P < 0.0001;* compared high glucose condition to the untreated condition (Vehicle), and SBI477 compared to high glucose condition (Mann Whitney U test). **(L-M)** Relative mRNA levels of *mlxipl* and *gcg* (normalized to *TBP*) in GLUTag cells incubated with 5 (G5) or 25mM glucose (G25) and transfected 48h with non-specific siRNA (ns) or siRNA targeting ChREBP (siChREBP) (n = 7-9). Values are means ± SEM. *\*P < 0.05; **P < 0.01*; compared each condition to ns (Mann Whitney U test). **(N)** Relative ChoRE-luciferase and *Gcg*-luciferase reporter activity in GLUTag cell line incubated with 5 (G5) or 25mM glucose (G25) and transfected with non-specific siRNA (ns) or siRNA targeting ChREBP (siChREBP). The Renilla luciferase reporter pRL-CMV was used as internal control. Dual luciferase reporter assays were performed 48h post transfection. Values are means ± SEM (n = 4). *\*P < 0.05; **P < 0.01*; compared G25 ns to G5 ns, and G25 siChREBP compared to G25 ns (Mann–Whitney U test).

### Ch^ΔGUT^ mice display delayed intestinal absorption of luminal glucose

Because intestinal glucose absorption is a key determinant of early glycemic control upon OGTT, we next evaluated functionally whether trans-epithelial glucose transport was altered in Ch^ΔGUT^ mice. Interestingly, the glycemic slope between 0 and 5 minutes after glucose gavage, which reflects the rate of intestinal glucose release into the bloodstream, was reduced by almost 35% in inducible Ch^ΔGUT^ mice when compared to CTL mice (Figure 4A). Similar results were observed in Ch*^−/−^* (Figure S5A) despite no difference in gastric emptying between Ch*^−/−^* and Ch*^+/+^* mice (Figure S2J). Since this suggested glucose malabsorption in Ch^ΔGUT^ mice, we recorded, though PET-Scan imaging over a 60-min period, the dynamic biodistribution of orally given 2-FDG tracer in CTL and Ch^ΔGUT^ mice. Slopes of bladder 2-FDG accumulation were used as an index of 2-FDG appearance in the bloodstream due to the lack of 2-FDG reabsorption by the kidneys. As illustrated in Figure 4B at 45min after the 2-FDG bolus, we noticed that 2-FDG appearance was significantly slower in bladder and in kidney from Ch^ΔGUT^ mice compared to CTL mice (Figure 4C-D). Interestingly, detection of 2-FDG signal was also delayed in liver, muscle, heart, lung and brain upon intestinal ChREBP deficiency (Figure 4E-I). Indeed, a significant delay was calculated to achieve 50% of maximal 2-FDG incorporation in the heart (30 min), the brain (13 min), the liver (10 min), the muscle (10 min), the lung (10 min), the bladder (10 min) and the kidney (8 min) of Ch^ΔGUT^ mice compared to CTL mice (Figure 4C-I). No difference was observed in the stomach between the 2 groups of mice, suggesting that slower biodistribution of 2-FDG to peripheral tissues was not attributable to a defect in gastric emptying (Figure 4J). Despite 2-FDG gut accumulation slopes revealed *in vivo* no significant increase upon gut ChREBP deficiency, 50% of end point 2-FDG signal was reached 10 min later in the gut of CTL mice as compared to Ch^ΔGUT^ mice (Figure 4K). Moreover, *ex vivo* measurement of 2FDG signal at 60 min after 2-FDG gavage highlighted a significant 1.85 fold raise in small intestinal 2-FDG contents of Ch^ΔGUT^ mice (Figure 4L-M), which was characterized by enhanced 2FDG accumulation in the proximal intestinal mucosa of Ch^ΔGUT^ mice (Figure 4N). Accordingly, measurement of transepithelial flux of ^14^C glucose in jejunal loop isolated from Ch*^−/−^* mice indicated that apical to basolateral glucose flux was significantly reduced upon ChREBP deficiency, to a similar extend of phloretin treated Ch*^+/+^* jejunum (Figure S5B). Altogether, these data pinpoint that, by impairing intestinal glucose absorption, loss of ChREBP activity in the gut epithelium may contribute to improved tolerance to oral glucose (Figure 2F).

**Figure 4:**
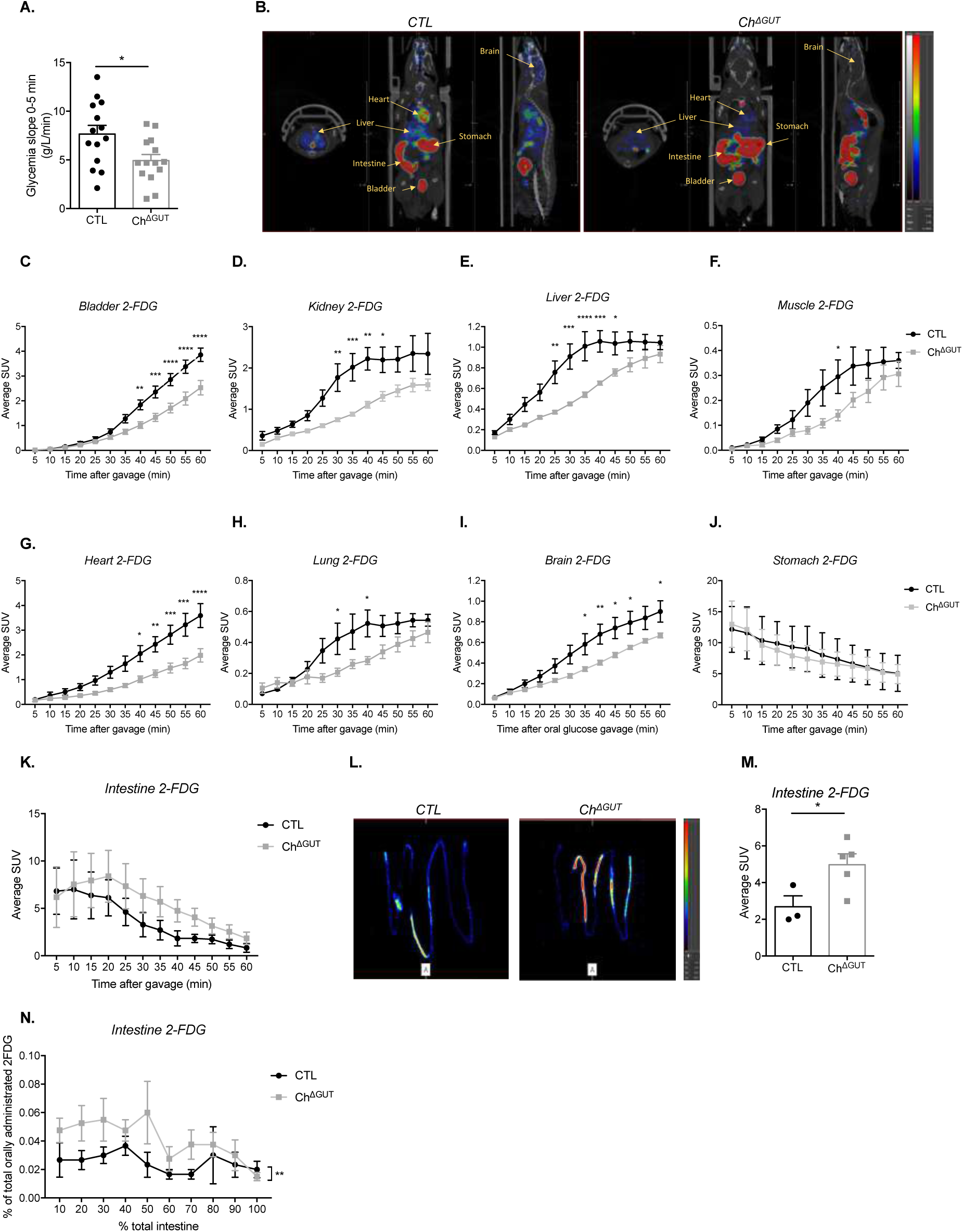
ChREBP^ΔGUT^ mice display delayed intestinal glucose absorption. **(A)** Glucose absorption index as measured by the glycemic slope between 0 and 5 min after oral glucose gavage (4g/kg) in 16-week-old male CTL and Ch^ΔGUT^ mice (n = 14). Values are means ± SEM. *\*P < 0.05*; compared CTL *vs* Ch^ΔGUT^ mice (Mann–Whitney U tests). **(B)** Representative transversal, coronal, and sagittal PET-Scan images of 2-FDG accumulation in 16-week-old male CTL and Ch^ΔGUT^ mice, 40 min after gavage of the tracer. **(C-K)** *In vivo* kinetic measure (over 60-min) of 2-FDG activity detected in **(C)** the bladder, **(D)** the kidney, **(E)** the liver, **(F)** the muscle, **(G)** the heart, **(H)** the lung, **(I)** the brain, **(J)** the stomach and **(K)** the intestine in 16-week-old male CTL and Ch^ΔGUT^ mice (n=4-5). Values are expressed as mean tissue radioactive concentration normalized by total orally administrated 2-FDG and body weight ± SEM (SUV: standardized uptake values). *\*P < 0.05, **P < 0.01, ***P < 0.001, ****P < 0.0001*; compared CTL *vs* Ch^ΔGUT^ mice two-way ANOVA followed by Bonferroni correction for multiple comparisons). **(L)** Representative PET-Scan images of 2-FDG accumulation in *ex vivo* flushed small intestine of 16-week-old male CTL and Ch^ΔGUT^ mice, 40 min after gavage of the tracer. PET figures and subsequent ones displayed intensity scale for tracer activity, from red (highest), trough green (intermediate) to blue (lowest). **(M)** *Ex vivo* tissue profiling for 2-FDG contents of non-flushed small intestine of 16-week-old male CTL and Ch^ΔGUT^ mice, 60 min after gavage of the tracer. Values are expressed as mean tissue radioactive concentration normalized by total orally administrated 2-FDG and body weight ± SEM (SUV: standardized uptake values) (n = 3-5). *\*P < 0.05*; compared CTL *vs* Ch^ΔGUT^ mice (two-way ANOVA followed by Bonferroni correction for multiple comparisons). **(N)** *Ex vivo* tissue profiling for 2-FDG contents along the flushed small intestine of of 16-week-old male CTL and Ch^ΔGUT^ mice, 60 min after gavage with the tracer. Values are means ± SEM. *\*\*P < 0.01*; compared CTL *vs* Ch^ΔGUT^ mice (Two-way ANOVA).

### ChREBP activity in the gut orchestrates disaccharides digestion and monosaccharides transport

Because the brush border morphology of enterocytes closely correlates with the absorptive capacity of the small intestine, we examined the ultrastructure of jejunum microvilli by transmission electron microscopy (Figure 5A). Ch^ΔGUT^ mice exhibited a drastic shortening (35%) of brush border microvilli in the jejunum, associated with a discrete reduction of microvilli density (10%) as compared to CTL mice (Figure 5B-C), thereby mimicking a condition of food deprivation. In Ch*^−/−^* mice, only microvilli length was also significantly reduced when compared to Ch*^+/+^* mice (Figure S6A-C). Surprisingly, concentrations of plasma β-hydroxybutyrate, an alternative energy source during nutrient deprivation, were not modified in random fed and fasted Ch^ΔGUT^ mice compared to CTL mice (Figure 5D) as well as in Ch*^−/−^* compared to Ch*^+/+^* mice (Figure S6D), suggesting no compensation of impaired intestinal glucose absorption by ketone bodies production upon ChREBP deficiency. To gain further insight into the molecular mechanisms underlying ameliorated glycemic control of Ch^ΔGUT^ upon OGTT, we performed comparative transcriptomic analyses on jejunum epithelial cells isolated from Ch^ΔGUT^ and CLT mice following an overnight 20%-glucose challenge in drinking water. Figure 5E shows that 5 clusters of genes that were differentially regulated between CTL mice receiving water or glucose and Ch^ΔGUT^ mice challenged with glucose, among which clusters 1 & 2 were glucose induced genes downregulated upon intestinal ChREBP deficiency (Figure 5E). A total of 791 genes was upregulated by glucose as compared to water in CTL mice (fold change ≥ 1,3, *P value* < 0,01) but only 21 glucose-controlled genes significantly depended on functional ChREBP activity (fold change ≤ −1,3, *P value* < 0,01) (Figure 5F, Table 2). Among the 52 down-regulated genes in Ch^ΔGUT^ compared to CLT mice in response to glucose (fold change ≤ −1,3, *P value* ≤ 0,01), *mlxipl* was the most downregulated gene (Table 3), as confirmed by RT-qPCR analysis (Figure S6A). Further gene ontology analysis of the 52 down-regulated genes in Ch^ΔGUT^ demonstrated that intestinal glucosidase, hydrolase, catalytic, and transporter activities were affected by ChREBP loss of function (Figure 5G-I). Moreover, among biological processes and pathways, carbohydrate metabolism, energy pathway and transport were the most altered in the gut of Ch^ΔGUT^ mice in response to 20%-glucose challenge (Figure 5G-I). Of note, intestinal ChREBP deficiency was associated with decreased expression of genes encoding (i) carbohydrate hydrolytic enzymes: *gaa* (−1.65 fold vs CTL, *P* = 0.0031), *lct* (−1.37 fold, *P* = 0.13), *mgam* (−1.3 fold, *P* = 0.0118), *sis* (−1.85 fold, *P* = 0.0019), (ii) hexose transporters: *slc2a2* (−2.74 fold, *P* = 0.0028), *slc2a5* (−2.26 fold, *P* = 0.0614), *slc2a7* (−1.49 fold, *P* = 0.4678), *slc5a1* (−1.45 fold, *P* = 0.0281), (iii) glycolytic enzymes: *eno1* (−1.38 fold, *P* = 0.0041), *pklr* (−2.07 fold, *P* = 0.0828), (iv) fructolytic enzymes: *aldoA* (−1.5 fold, *P* = 0.1081), *aldoB* (−1.2 fold, *P* = 0.0051), *khk* (−1.69 fold, *P* = 0.0477)*, pfkl* (−1.36 fold, *P* = 0.7504), *(v)* galactolytic enzymes: *galk2* (−1.24 fold, *P* = 0.1210), and (vi) neoglucogenic enzyme: *g6pc* (−1.31 fold, *P* = 0.1148) (Figure 5J). This suggested that intestinal ChREBP activity controls a transcriptional program orchestrating efficient sugar digestion, transport and metabolism. Accordingly, RT-qPCR analyses on jejunal epithelial cells isolated from Ch^ΔGUT^ and Ch*^−/−^* mice validated that genes encoding hexose transporters (*slc2a2*, *slc2a5*, *slc2a7, slc5a1*) require ChREBP activity (Figure 5K, S6F). Moreover, mRNA levels of disaccharidase (*sis, lct, mgam*) or hexose metabolism (*galk2*, *g6pc, khk, lpk, tkfc*) were significantly reduced in the gut mucosa of Ch^ΔGUT^ or in Ch*^−/−^* mice as compared to CTL or Ch*^+/+^* mice (Figure 5K, S6G). Of note, intestinal expression of brush border markers (*pept1, fabp, cd36*), which encode di/tri-peptide and lipid transporter respectively, was investigated in Ch^ΔGUT^ or Ch*^−/−^* mice (Figure 5L, S6G), and results suggest a specific role of ChREBP in brush border sugar hydrolysis and absorption despite reduced microvilli length in both mice models (Figure 5B, S6C). Collectively, these results emphasized that in addition of hampering sugar transport across the gut epithelium, abrogation of the transcriptional program driven by intestinal ChREBP impairs efficient luminal digestion of carbohydrates.

**Figure 5:**
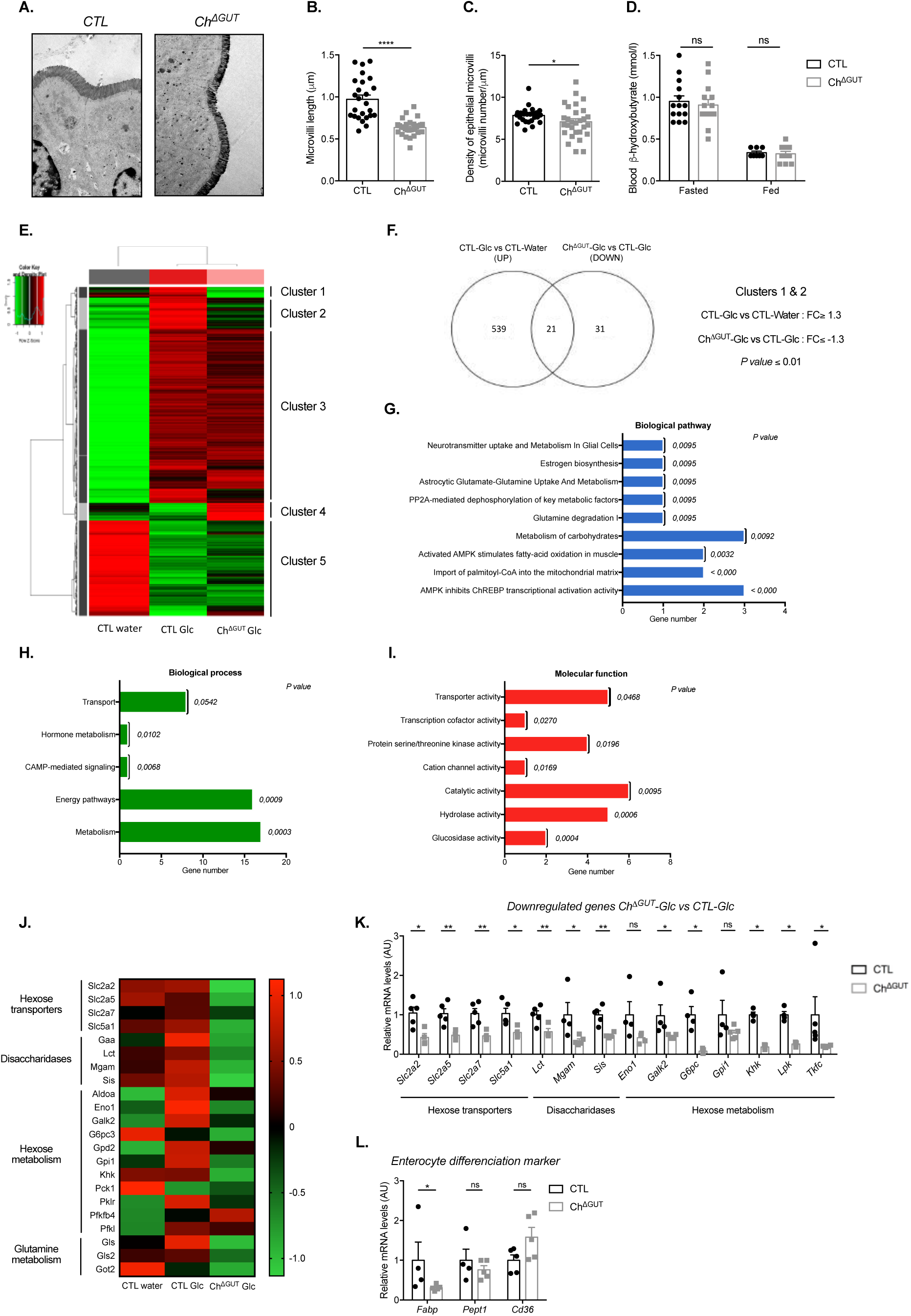
ChREBP activity controls intestinal expression of disaccharidases and hexose transporters. **(A)** Representative transmission electron photomicrographs of the jejunal brush border from 16-week-old random fed male CTL and Ch^ΔGUT^ mice (Magnification ×2000). Scale bar: 1µm. **(B-C)** Length and density of microvilli in the jejunum of 16-week-old random fed male CTL and Ch^ΔGUT^ mice (n = 3). **(D)** Plasma β-hydroxybutyrate levels in fasted and fed 16-week-old male CTL and Ch^ΔGUT^ mice (n = 8-14). **(B-D)** Values are means ± SEM. *\*P < 0.05*, *\*\*\*\*P < 0.0001*; compared CTL *vs* Ch^ΔGUT^ mice (Mann Whitney U test). **(E-J)** Transcriptomic analyses performed on jejunum epithelial cells isolated from Ch^ΔGUT^ and CLT mice after an overnight with 20%-glucose challenge (Ch^ΔGUT^ Glc, CTL Glc) in drinking water and compared to untreated CTL mice (CTL Water) (n = 4). **(E)** General heatmap of differentially expressed probes. Color bar indicates the range of expression levels in the heatmap (green: downregulation red: upregulation). **(F)** Venn diagram presenting the overlap between genes upregulated in jejunal mucosa of CTL Glc mice compared to CTL Water (fold change ≥ 1,3; *P value* < 0,01) and genes downregulated in Ch^ΔGUT^ Glc mice compared to CTL Glc mice (fold change ≤ −1,3; *P value* < 0,01) performed using Venny^2.1^. **(G-I)** Gene Ontology (GO) analysis of genes downregulated in jejunal mucosa of Ch^ΔGUT^ Glc mice compared to CTL Glc mice generated through IPA and FunRich softwares (fold change ≥ 1,3; *P value* < 0,01). The x-axis represents the number of differentially expressed genes based on the dataset from FunRich software while the y-axis corresponds at the annotation of molecular function. *P value* (≤0,05) are included in the bar graph. **(J)** Selected heatmap of genes related to sugar digestion, transport and metabolism in jejunal mucosa of CTL water, CTL Glc and Ch^ΔGUT^ Glc mice. Color bar indicates the range of expression levels in the heatmap (green: downregulation red: upregulation). Abbreviations: *slc2a2* for GLUT-2; *slc2a5* for GLUT-5; *slc2a7* for GLUT-7; *slc5a1* for SGLT-1; *gaa* for Maltase; *lctl* for Lactase; *mgam* for Maltase-Glucoamylase; *sis* for Sucrase Isomaltase; *aldoA* for Aldolase A; *eno1* for Enolase 1; *galk2* for Galactokinase 2; *g6pc3* for G6pase 3; *gpd2* for Glycerol-3-Phosphate Dehydrogenase 2; *gpi1* for Glucose-6-phosphate isomerase; *khk* for Ketohexokinase; *pck1* for Phosphoenolpyruvate carboxykinase; *Pklr* for Pyruvate Kinase; *Pfkfb4* for 6-Phosphofructo-2-kinase/fructose-2,6-biphosphate 4; *pfkl* for 6-phosphofructokinase; *gls* for Glutaminase; *gls2* for Glutaminase 2; *got2* for Aspartate aminotransferase. **(K, L)** Relative mRNA levels (normalized against *TBP*) in jejunal epithelial cells from 16-week-old male CTL and Ch^ΔGUT^ mice (n = 4-5). Values are means ± SEM. *ns, *P < 0.05*, *\*\*P < 0.01;* compared CTL *vs* Ch^ΔGUT^ mice (Mann– Whitney U tests).

**Table 2:**
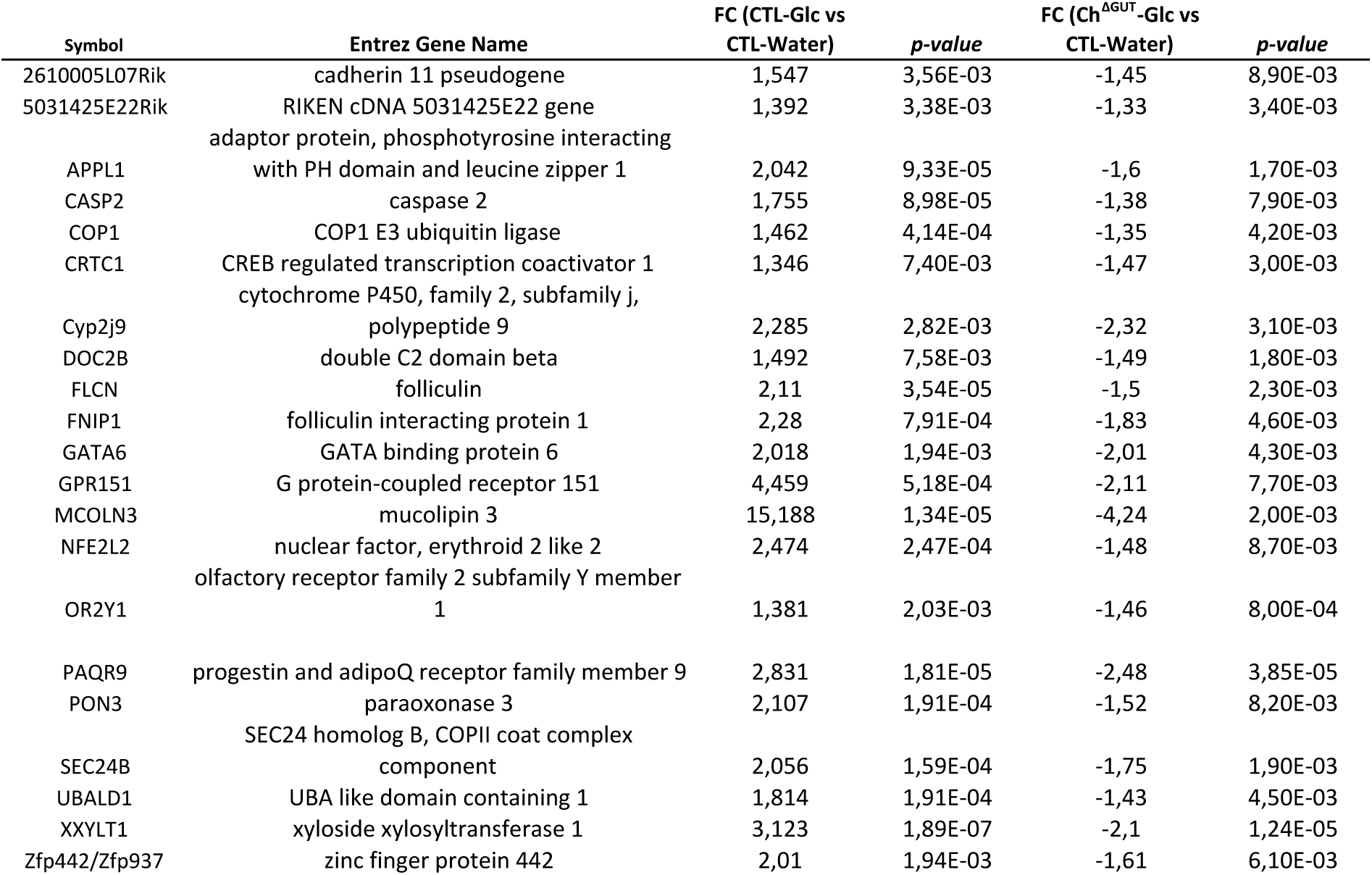
Transcriptomic analysis of the 21 glucose-induced genes (fold change (FC) ≥ 1,3 with *p-value* < 0,01) down-regulated upon intestinal ChREBP deficiency (fold change (FC) ≤ - 1,3 with *p-value* < 0,01) in jejunal epithelial cells. Fold changes and *p-values* detailed below compare Ch^ΔGUT^ mice versus CTRL mice, both upon overnight 20% glucose challenge in the drinking water

**Table 3:**
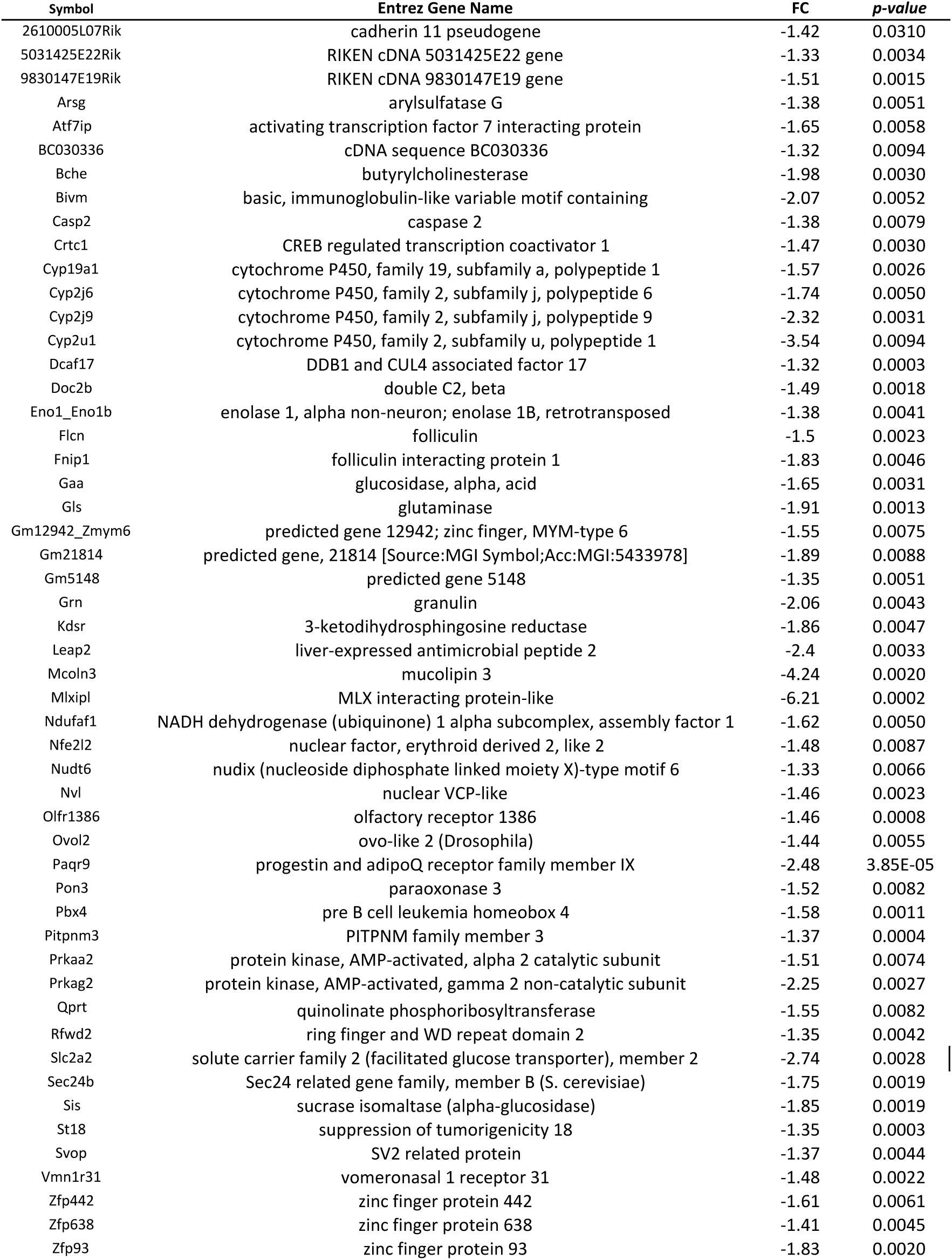
Transcriptomic analysis of the 52 genes down-regulated upon intestinal ChREBP deficiency (fold change (FC) ≤ −1,3 with *p-value* < 0,01) in jejunal epithelial cells. Fold changes and *p-value* detailed below compare Ch^ΔGUT^ mice versus CTRL mice, both upon overnight 20% glucose challenge in the drinking water.

### Intestinal ChREBP deficiency induces early intolerance to high-sucrose and high-lactose diets

Because gene expression of major disaccharidases (sucrase-isomaltase, lactase) was reduced upon intestinal ChREBP deficiency, we next compared the short-term (4 days) and long-term (40 days) consequences of high-sucrose or high-lactose feeding in Ch^ΔGUT^ and CTL mice. As shown in Figure 6A, macroscopic examination of gut from Ch^ΔGUT^ mice fed a 60% sucrose diet revealed exacerbated cecal and intestinal distention, as characterized by enhanced gas and liquid contents, when compared to CTL mice. Accordingly, caecum weight was increased in Ch^ΔGUT^ mice after both 4 days (2.9 fold vs CTL) or 40 days (1.4 fold vs CTL) of high-sucrose feeding, despite similar weight loss in both groups of mice (Figure 6A-C), thereby suggesting enhanced sucrose intolerance upon intestinal ChREBP deficiency. Of note, higher cecal distension was observed in Ch*^−/−^* mice fed a high-sucrose during a short time frame, as the caecum weight was increased by 5.5 fold compared to Ch*^+/+^* mice (Figure S7A). Compatible with intolerance to sucrose upon gut ChREBP loss, the mean daily food intake was reduced by 43% in Ch^ΔGUT^ mice fed for 4 days with a 60% sucrose diet as compared to the CTL group (Figure 6D). Hyperglycemia and enhanced liver weight induced by a 4 days high-sucrose diet exposure in CTL mice, were totally reversed Ch^ΔGUT^ mice (Figure 6E-F). Analysis of the cecal flora from both groups of mice highlighted that, following short term exposure with high-sucrose, intestinal ChREBP loss of function resulted in higher abundance of the *Actinobacteria* phylum, in particular *Bifidobacterium spp.* (Figure 6G-H). In agreement with previous study on global ChREBP knock-out mice (Kato et al., 2018), this suggested that higher amount of unabsorbed sugars in the distal gut lumen of Ch^ΔGUT^ mice fed a high-sucrose diet favored the growth of bacterial species which can degrade hexose sugars (Devika & Raman, 2019; Pokusaeva et al., 2011). No significant change in *Firmicutes* and *Proteobacteria* phyla and *Bacteroides spp.* was observed in Ch^ΔGUT^ mice fed a 60% sucrose diet (Figure 6I-J). While exposure to 60% lactose for 4 days resulted in an early intolerance to the diet in both CTL and Ch^ΔGUT^ mice, this response was however aggravated Ch^ΔGUT^ mice, as illustrated by enhanced caecum weight (a 2.3 fold increase vs CTL) and body weight loss (1.6 fold vs CTL) after 4 days of diet (Figure 6A-C). Similarly, evaluation of cecal distension in Ch*^−/−^* mice fed a high-lactose diet during a short time frame revealed even more pronounced effects, as the caecum weight was increased by 2.3 fold when compared to Ch*^+/+^* mice (Figure S7A). Of note, no difference in cecal or body weight was observed between Ch^ΔGUT^ and CTL mice at 40 days of high-lactose challenge, likely due to the concomitant onset of massive lactose intolerance in CTL mice (Figure 6A-C) and consistent with the weak expression of lactase in adult mice (Seetharam et al., 1977). However, examination of bacterial load in the caecum revealed that Ch^ΔGUT^ mice exhibit a lower cecal abundance of the *Bacteroides* genus and *Prevotella spp* compared to CTL mice with no change in *Bifidobacterium spp* (Figure 6H-J). Of note, we reported no lethality in Ch^ΔGUT^ or Ch*^−/−^* mice fed with either a high-sucrose diet or a high-lactose diet. Altogether, these data indicate that intestinal ChREBP deficiency causes an early intolerance to high-disaccharide diets such as sucrose and lactose.

**Figure 6:**
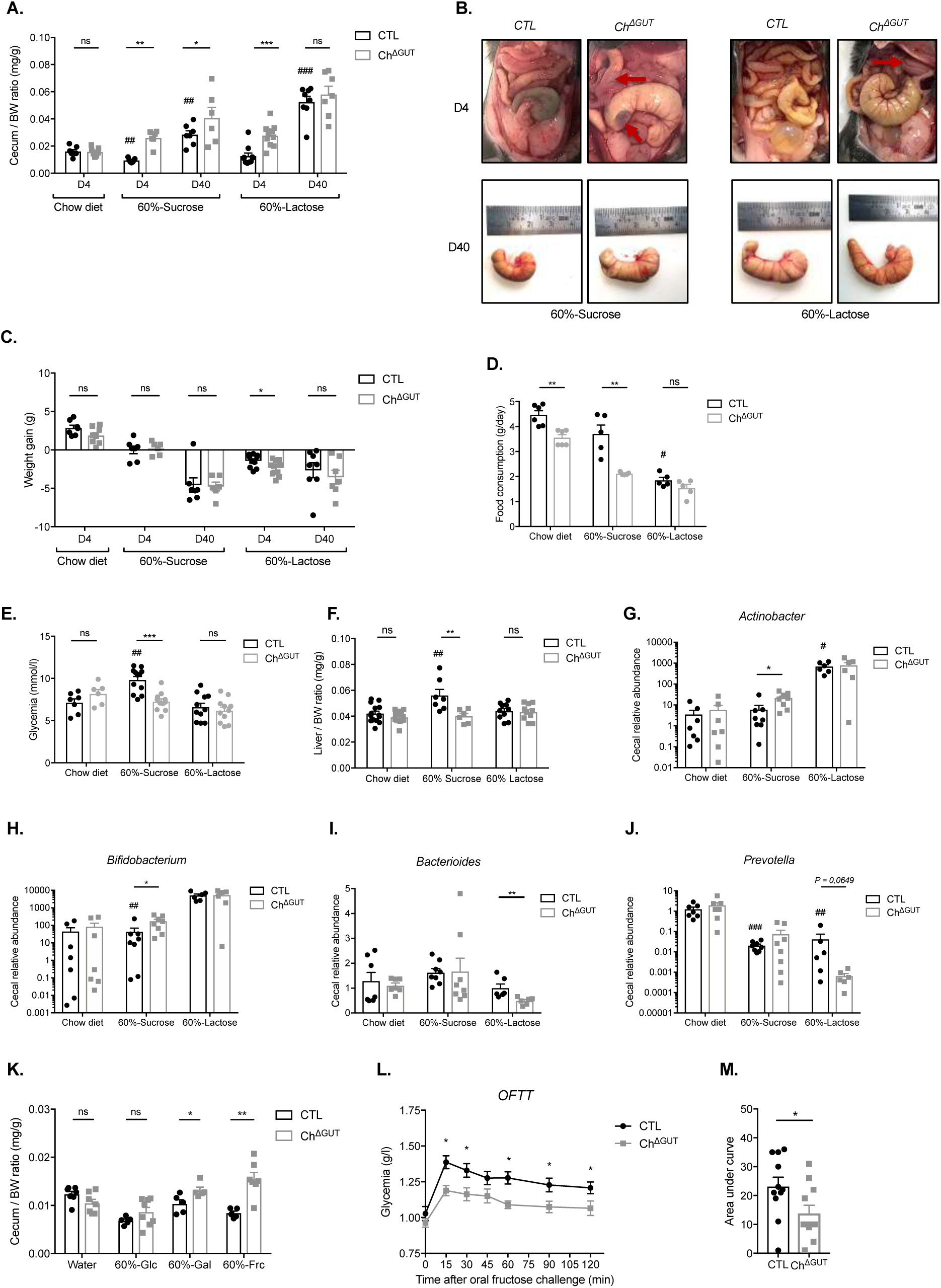
Intestinal ChREBP deficiency induces early intolerance to high sucrose and lactose as well as malabsorption of fructose and galactose. **(A-J)** 16-week-old male CTL and Ch^ΔGUT^ mice fed either with 60% sucrose or 60% lactose diets for 4 four (D4) or forty days (D40). (A) Representative images of cecum *in situ* or *ex vivo*. **(B)** Relative caecum weight (normalized to body weight), **(C)** body weight gain (n = 6-10), **(D)** daily food consumption per mice (over a 4 days period) (n = 5-6) and **(E)** plasma glucose (after 4 days of diet) (n = 5) and **(F)** relative liver weight (normalized to body weight) (after 4 days of diet) (n = 7-14). **(G-J)** Relative abundance of *Actinobacter, Bifidobacterium, Bacteroides and revotella* in cecal feces collected form mice after 40 days of diet (n = 6-8). Values were normalized against *UTB.* **(A-J)** Values are means ± SEM. *^#^P < 0.05, ^##^P < 0.01;* compared CTL sucrose/lactose diet *vs* chow diet (Mann–Whitney U test). *ns, *P < 0.05, **P < 0.01, ***P < 0.001*; compared CTL *vs* Ch^ΔGUT^ mice (Mann Whitney U test). **(K)** Relative caecum weight (normalized to body weight) in 16-week-old male CTL and Ch^ΔGUT^ mice which received either water or 60%-glucose, 60%-fructose or 60%-galactose in the drinking water for 3 days. Values are means ± SEM (n = 5-8). *ns, **P < 0.01*; compared CTL *vs* Ch^ΔGUT^ mice (Mann–Whitney U test). **(L)** Kinetic measurement of blood glucose and **(M)** corresponding area under curves (AUC) upon an oral load of fructose (2g/kg) performed in 16-week-old male CTL and Ch^ΔGUT^ mice after an overnight fasting (n = 11). Values are means ± SEM. *\*P < 0.05*; compared control *vs* Ch^ΔGUT^ mice (Mann Whitney U test).

### Intestinal ChREBP deficiency triggers both galactose and fructose malabsorption syndroms

We next investigated whether, along with impaired breakdown of disaccharides, monosaccharide malabsorption could participate to the onset of sucrose and lactose intolerance upon intestinal ChREBP deficiency. The relative contribution to this phenotype of the three hexoses, that make up sucrose (glucose-fructose) and lactose (glucose-galactose), was assessed by challenging Ch^ΔGUT^ and CTL mice with 60% glucose or 60% fructose in drinking water for 3 days (Figure 6L). Because a 60% galactose challenge in drinking water induced lethality in both CTL and Ch^ΔGUT^ mice (data not shown), we performed daily gavages with a 60% galactose solution (3 gavages/day, 1% BW). While we noticed no statistical change in caecum weight of Ch^ΔGUT^ mice compared to the CTL group upon a high-glucose challenge, cecal contents were significantly increased in Ch^ΔGUT^ compared to CTL mice in response to both high-galactose and high-fructose exposure (Figure 6K), indicating that intestinal ChREBP deficiency induced galactose as well as fructose malabsorption syndromes. Of note, the intolerance to fructose was more pronounced than that to galactose, as illustrated by the 1.3 and 1.9 fold increase in caecum weight in Ch^ΔGUT^ respectively when compared to the CTL group. The severe fructose intolerance of Ch^ΔGUT^ mice was not associated with any lethality, in contrast to previous observations in mice with a constitutive and total deletion of *mlxipl* in the gut (Kim et al., 2017). Because the small intestinal epithelium was recently identified as the main site of dietary fructose clearance in mice (Jang et al., 2018), we next investigated whether intestinal ChREBP deficiency could impact glycemic excursion upon an acute fructose oral challenge. As shown in Figures 6L and 6M, fructose gavage indeed reduced by 40.2% decrease of glycemic excursion in Ch^ΔGUT^ mice as compared to the CTL group, indicating that intestinal ChREBP activity is essential for local fructose absorption and metabolism and subsequent elevation of blood glucose concentrations.

## DISCUSSION

In the current study, the contribution of the intestinal transcription factor ChREBP to glycemic control was evaluated *in vivo* using both total *mlxipl* knock out mice (Ch^-/-^) and mice with a targeted inducible deletion of *mlxipl* specifically in the gut epithelium (Ch^ΔGUT^). We demonstrate here that improved tolerance to oral glucose challenge upon intestinal ChREBP deficiency is accompanied by a paradoxical decrease of incretin response but impaired transepithelial glucose absorption. Among potential molecular mechanisms underlying intestinal ChREBP functions, we revealed that ChREBP orchestrates the local expression of several hexose transporters as well as disaccharidases. Consequently, ChREBP gut deficiency triggers an early intolerance to dietary lactose and sucrose, mainly due to galactose and fructose malabsorption.

Our data demonstrate that ameliorated tolerance to an oral glucose load upon intestinal ChREBP deficiency was not accompanied by enhanced secretion of incretin hormones, but rather by a reduction of glucose-induced GLP-1 secretion and delayed intestinal glucose absorption. Confirming previous observations (Habib et al., 2012; Trabelsi et al., 2015), we highlight that ChREBP is enriched in intestinal GLP-1 producing L-cells. While specific expression of glucokinase was shown to control intracellular glucose metabolic flux in L cells (Parker et al., 2012; Reimann et al., 2008), this suggests that, like in hepatocytes, ChREBP could be activated by glucose-derived metabolites. We report that loss of ChREBP activity in the intestine induces a potent downregulation of epithelial mRNA levels encoding the Na^+^-coupled glucose transporter SGLT1, which was previously established as the major glucose-sensing mechanism triggering GLP-1 secretion *in vitro* in primary L cells (Parker et al., 2012), *ex vivo* in perfused intestinal loops (Moriya et al., 2009) and *in vivo* in SGLT-1 knockout mice (Gorboulev et al., 2012). Moreover, the high expression level of SGLT1 in the proximal small intestine (Yoshikawa et al., 2011) indicates overlapping expression patterns of SGLT1 and ChREBP. Therefore, the early decreased portal GLP-1 concentrations in Ch^ΔGUT^ mice in response to luminal glucose challenge could result from impaired electrogenic glucose uptake though SGLT-1 in the gut epithelium. Accordingly, SGLT-1 mRNA levels were significantly reduced in L-cells from Ch^ΔGUT^ mice as well as in GLUTag cells upon ChREBP knock down (data not shown). Of note, *Oh A et al.* previously reported no significant change in intestinal SGLT-1 expression under basal or oral glucose challenge but reduced *slc5a1* mRNA levels following acute or chronic exposure to dietary fructose upon total ChREBP deficiency (Oh et al., 2018).

Besides its potential contribution to the regulation of L cells granules exocytosis, we report in this study that gut epithelial *gcg* expression is reduced in Ch^ΔGUT^ mice and that the genetic or pharmacological inhibition of ChREBP in GLUTag cells prevents the increase in *gcg* mRNA levels following high glucose challenge. Glucose-induced proglucagon gene expression was previously shown to rely on glucose metabolism and FXR interaction with ChREBP was reported to inhibit glucose-induced *gcg* expression in the GLUTag cell lines (Trabelsi et al., 2015). Taken together, these data indicate that, beyond GLP-1, ChREBP could regulate the production of other proglucagon-derived enteropeptides, such as GLP-2, glicentin or oxyntomodulin. While our results suggest that ChREBP could transactivate *gcg* promotor, bioinformatic analysis did not reveal ChREBP binding site in the *gcg* promoter (Poungvarin et al., 2015) (data not shown), putative ChoRE consensus sequences were identified *in silico* at −45 and −259 of the *pax6* promoter (data not shown). Pax-6 was demonstrated to bind to *gcg* promoter, and mice homozygous for a dominant negative *pax6* allele exhibit reduced intestinal proglucagon mRNA levels as well as undetectable GLP-1-immunopositive cells in the gut mucosa (Hill et al., 1999; Trinh et al., 2003). Of note, *pax6* expression is induced by glucose in rat insulinoma INS-1E cells (Balakrishnan et al., 2014) and *mlxipl* expression was shown to parallel those of *pax6* and *gcg* upon L-cells adaptation to high fat diet (Dusaulcy et al., 2016). Moreover, our data evidence that *pax6* expression is significantly decreased upon ChREBP knock down in GLUTag cells as well as in the intestinal mucosa of Ch^ΔGUT^ mice (data not shown). Furthermore, L cells number was reduced in small intestinal mucosa of Ch^-/-^ mice, with no alteration in colonic L cells, consistent with ChREBP distribution pattern along the gut and with no significant enrichment in ChREBP expression in colonic L cells. Further investigations will be needed to determine whether ChREBP binds directly on *gcg* promoter or whether it could indirectly stimulate *gcg* transcription through Pax6, thereby affecting L cell differentiation.

Dynamic 2-FDG microPET scan analyses revealed that Ch^ΔGUT^ mice display intestinal glucose malabsorption and delayed glucose uptake by peripheral tissues, with no parallel modification of gastric emptying, a major determinant of glucose absorption kinetic (Heading, 1994). Mechanistically, we show that intestinal ChREBP activity controls the local expression of genes encoding SGLT-1, GLUT-2 and GLUT-7 glucose transporters (Cheeseman, 2008). While apical glucose uptake is predominantly mediated by SGLT1, its passive basolateral release from intestinal epithelial cells involves GLUT2 (Röder et al., 2014). However, adaptive absorption of glucose across the brush-border was reported upon post-prandial or insulin-resistant conditions through apical GLUT2 translocation (Affleck et al., 2003; Ait-Omar et al., 2011; Boudry et al., 2007; Chaudhry et al., 2012; Grefner et al., 2015; Zheng et al., 2012). Moreover, *slc5a1* expression was shown to be upregulated in the proximal gut of diabetic rodent models (Song et al., 2016) and the subsequent enhanced SGLT1-mediated intestinal glucose uptake was proposed to contribute to the rapid post-prandial rise in blood glucose levels observed in diabetes (Powell et al., 2013; Zambrowicz et al., 2012). Therefore, by controlling both *slc5a1* and *slc2a2* expression, ChREBP activity in the gut epithelium might represent a key player of intestinal glucose absorption and its inhibition might have beneficial effects on postprandial glycemic control upon diabetic conditions. Accordingly, we reveal an improved oral glucose tolerance in Ch^ΔGUT^ mice, as observed in SGLT-1 and GLUT2^ΔGUT^ deficient mice (Gorboulev et al., 2012; Schmitt et al., 2017). Consistent with our data, previous studies documented that reduced gut *slc5a1* and *slc2a2* mRNA levels upon total or constitutive intestinal deletion of ChREBP after feeding with sucrose or fructose (Kato et al., 2018; Kim et al., 2017; Oh et al., 2018). While decreased *slc2a2* expression was also reported in the liver of Ch^-/-^ mice (Iizuka et al., 2004), two independent ChIP sequencing analyses failed to identify *slc2a2* as a direct ChREBP target gene (Jeong et al., 2011; Poungvarin et al., 2015). Interestingly, the PPAR signaling pathway was identified among the top genes with high ChREBP binding peak intensity in both hepatocytes and white adipocytes (Poungvarin et al., 2015) and heterodimers of PPARα or PPARγ and RXRα were shown to stimulate *slc2a2* expression through direct binding to PPRE promoter sequences (Cha et al., 2000; Im et al., 2005). Our transcriptomic analyses revealed that genes endoding PPARγ and PGC1α were downregulated by 1.4 and 2.7 fold upon intestinal ChREBP deficiency (data not shown). Further investigations will be needed to determine whether ChREBP directly or indirectly controls *slc5a1, slc2a2* and *slc2a7* transcription.

Interestingly, we report that lowering effects of intestinal ChREBP deficiency on post-absorptive glycemia is not restricted to glucose consumption but also occurs upon fructose challenge. Recent evidence demonstrate that most of low-dose luminal fructose appears in the portal blood as glucose and lactate, suggesting exposure of the liver to a similar metabolic milieu as does glucose ingestion (Jang et al., 2018). In this context, we demonstrate that inducible deletion of *mlxipl* in the gut prevents the local expression of genes encoding fructose transporters (*slc2a2, slc2a5)*, fructolytic (*khk, aldoA*) and gluconeogenic (*g6pc*) enzymes, which are required for glucose production from fructose. Accordingly *slc2a5* was demonstrated as a direct ChREBP target gene in human Caco-2 cells and in mouse jejunal mucosa (Kim et al., 2017; Oh et al., 2018) and two ChoRE regions were identified in the human *khk* promoter (Lanaspa et al., 2012). Considering that *slc2a5* and *khk* gene deletion both suppress fructose-induced intestinal gene expression of *mlxipl* and *lpk*, a ChREBP prototypic target gene (data not shown), fructose-derived metabolites might regulate fructolytic gene expression through ChREBP activation. Interestingly, several studies indicated that increased intestinal absorption of fructose stimulates local lipid packaging into chylomicrons (Egli et al., 2013; Haidari et al., 2002; Theytaz et al., 2014), suggesting that by curbing intestinal fructose uptake and metabolism, gut ChREBP loss may contribute to reduce fructose-induced dyslipidemia. Moreover, because high doses of fructose were reported to overwhelm fructose metabolic capacities of the small intestine and to spill over to the liver where it stimulates *de novo* lipogenesis (Jang et al., 2018), it is tempting to speculate that upon high-sucrose or -fructose dietary intake, the blockade of intestinal ChREBP-mediated *slc2a5* expression might have beneficial outcomes on hepatic steatosis by curbing both glucose and fructose portal fluxes. Deletion of *slc2a5* in mice fed a high fructose diet indeed resulted in a 90% decrease of serum fructose levels (Barone et al., 2009). However, improvement of fructose-induced liver steatosis upon intestinal ChREBP deficiency remains uncertain as our results indicate higher abundance of acetate-producing *Bifidobacterium* species in the caecum of Ch^ΔGUT^ mice. While conversion by the gut microbiota of dietary fructose to acetate was previously reported (Jang et al., 2018), microbial derived acetate was shown to feed hepatic lipogenesis (Zhao et al., 2020). Therefore, increased production of microbial acetate upon intestinal ChREBP deficiency may contribute to a raise of lipogenic acetyl-CoA pools in the liver. Consistent with the abrogation of sucrose-enhanced liver weight in Ch^ΔGUT^ mice, Kim et al suggested that the reduction of fructose-mediated accumulation of hepatic cholesterol and triglycerides upon constitutive deletion of ChREBP in the intestine likely results from fructose intolerance and consecutive impairment of food intake and weight loss (Kim et al., 2017).

Besides its implication in hexose absorption, we demonstrate that intestinal ChREBP activity controls gene expression of disaccharidases involved in the hydrolytic step of luminal dietary sugars. Consistent with this, significant diminution of *sis* mRNA levels was previously documented in proximal intestine of Ch^-/-^ mice fed a 30% sucrose diet (Kato et al., 2018). Of note, diabetic conditions have been reported to trigger the increase of disaccharidase activity (sucrase, isomaltase, maltase, lactase) in rodent experimental models and patients (Adachi et al., 1999; Ikeda, Murakami, 1998; Olsen & Korsmo, 1977; Tandon et al., 1975) and α-glucosidase inhibitors are used as an adjunctive type 2 diabetes therapy for their lowering effect on postprandial glycemia and insulin levels (Chiasson et al., 2002; Hanefeld, 2007; Inzucchi et al., 2015; Yang et al., 2014). Moreover, the elevation of disaccharidase activities upon diabetic state was suggested to be independent of enteral factors (luminal nutrients, gastrointestinal hormones) but rather to be directly regulated by hyperglycemia (Schedl et al., 1983). Further analyses will be necessary to determine whether hyperglycemia could stimulate ChREBP activity locally and thereby contribute to exacerbated disaccharides digestion and enhanced sugar absorption upon diabetic conditions.

While the targeting of intestinal ChREBP might display some anti-diabetic effects by hampering dietary sugars breakdown and hexose transport, the blockade of its activity could trigger adverse side effects in case of high-sugar diets. Of note, in the context of the treatment for diabetes, metformin was previously shown to reduce ChREBP activity in hepatocytes (Sato et al., 2016), INS-1E cells (Shaked et al., 2011) and endothelial cells (Li et al., 2015b). Therefore, while treatment with metformin is frequently associated with abdominal discomfort (diarrhea and vomiting) (Sanchez-Rangel & Inzucchi, 2017), parallel inhibition of intestinal ChREBP activity might contribute to abdominal side effects of metformin therapies. Indeed, we show that Ch^ΔGUT^ mice developed an early intolerance to high sucrose challenge due to fructose malabsorption, as characterized by bloating, diarrhea and cecal distension likely due to bacterial fermentation of unabsorbed fructose in the colon. While our results indicate that intestinal ChREBP activity controls the expression of *sis* and *slc2a5*, similar symptoms were observed in patients with congenital *Sis* deficiency or in *slc2a5^-/-^* mice fed a high fructose diet (Barone et al., 2009). In contrast to a previous study (Kim et al., 2017), we reported no lethality in high-fructose fed Ch^ΔGUT^ mice, likely due to different nutritional interventions, as mice had free access 60% fructose in drinking water and to chow diet in our study. Despite intestinal fructose intolerance was not associated with intestinal GLUT5 and GLUT2 mRNA and protein levels in patients (Wilder-Smith et al., 2014), future evaluation of gut ChREBP expression and activity might elucidate whether ChREBP-mediated transcription could be one of the mechanisms underlying human fructose malabsorption, which are still largely unknown. Moreover, the prevalence and severity of fructose malabsorption being directly proportional to dietary fructose levels and inversely proportional to age (Gomara et al., 2008; Jones, et al., 2011), further analysis would be of interest in order to determine whether ChREBP activity is mandatory to trigger *scl2a5* expression at the sucking-weaning transition (Davidson et al., 1992; Douard & Ferraris, 2013). In this context, our data indicate that intestinal expression of *mlxipl* and ChREBP prototypic target genes are significantly enhanced in weaned mice upon dietary change from high-fat milk to high-carbohydrate chow (Figure S8A).

Besides intolerance to high-sucrose and high-fructose diets, we report that Ch^ΔGUT^ mice exhibit aggravated intolerance to lactose, in parallel with reduced levels of *lct* mRNA. While this suggested that, upon ChREBP deficiency, impaired residual activity of lactase contributes at the adult stage to enhanced fermentation of indigestible dietary lactose in the distal gut by the microflora (Van de Heijning et al., 2015), we did not observe significant change in cecal abundance of the lactose-digesting *Bifidobacterium* species. Evaluation of cecal *Lactobacillus* relative abundance in Ch^ΔGUT^ mice will complement these data. Interestingly, the marked lactose intolerance upon loss of ChREBP activity was accompanied by intestinal malabsorption of galactose, consistent with reduced intestinal expression of SGLT-1 in Ch^ΔGUT^ mice. While dietary lactose is normally broken down into glucose and galactose by lactase, hexoses are further transported into intestinal epithelial cells by the Na+-glucose cotransporter SGLT1. Thus, *slc5a1* genetic defects are associated with intestinal malabsorption of glucose and galactose in humans (Martín et al., 1996a; Turk et al., 1991) and similar symptoms have been observed in mice lacking *slc5a1* (Gorboulev et al., 2012).

Finally, a poorly understood feature of the metabolic syndrome is its association with leaky gut, which is suggested to nurture a chronic low-grade inflammatory state (Winer et al., 2017). Hyperglycemic high-glucose or -fructose diets were shown to increase gut permeability subsequently to cell-cell junction alterations to the tight junction proteins (Do et al., 2018). In this context, *Thaiss et al*. recently demonstrated that hyperglycemia drives loss of epithelial integrity and higher risk for enteric infection by triggering a retrograde glucose flux into gut epithelial cells and by causing a transcriptional reprogramming subsequent to alterations of intracellular glucose metabolism (Thaiss et al., 2018). Our data show that mRNA levels of *mlxipl* and *lpk* are enhanced in jejunal epithelial cells of hyperglycemic *ob/ob* or streptozotocin-treated mice (Figure S8C-F), suggesting that hyperglycemia could stimulate intestinal ChREBP activity. While loss of ChREBP in mouse small bowel is sufficient to significantly lower *slc2a2* and *gcg* expression, hyperglycemia-induced intestinal ChREBP activity could deteriorate gut permeability through regulation of GLUT-2-mediated glucose metabolism and proglucagon-derived peptides production. Intestinal GLUT-2 deletion was indeed shown to restore barrier function and bacterial containment in a diabetic mice model despite sustained hyperglycemia (Thaiss et al., 2018). Moreover, pharmacological treatment with GLP-2 decreased gut permeability and systemic inflammation associated with obesity, whereas GLP-2 antagonist abolished most of prebiotic beneficial effects on gut barrier in *ob/ob* mice (Cani et al., 2009). Surprisingly, our data show that gut epithelial permeability is reduced by 40% in Ch^ΔGUT^ mice under basal conditions, as observed *ex vivo* upon intestinal deficiency on GLUT-2 (Schmitt et al., 2017). Therefore, investigating whether invalidation of ChREBP in the gut epithelium can reverse the intestinal hyperpermeability and ameliorate the systemic inflammation and the risk of enteric infections of diabetic mice models either fed a high-fat diet or treated with streptozotocin would be of great interest.

In summary, our study demonstrates that intestinal ChREBP activity orchestrates a transcriptional program, which is essential for the digestion of dietary sugars and transepithelial absorption and the intracellular metabolism of luminal monosaccharides. We provide new insights into the molecular mechanisms controlling sugar absorption and GLP-1 production in the intestinal epithelium by ChREBP. In a context of type 2 diabetes, targeting ChREBP may be a useful strategy to reduce postprandial glycemic excursion, as already used in anti-diabetic therapies (alpha-glucosidase inhibitors) but the blockade of intestinal ChREBP activity may be accompanied by an accumulation of undigested sugars in the colon thereby contributing to the development of sugar intolerance.

## ACKNOWLEGMENTS

We are grateful to Dr Daniel P Kelly (Center for Metabolic Origins of Disease, Sanford Burnham Prebys Medical Discovery Institute, Orlando, Florida, USA) for the kind gift of SBI-477, a ChREBP inhibitor. We also thank the RPPA mouse facility and Amélie Lacombe from the PreClinICAN metabolic phenotyping facility (Institute of Cardiometabolism and Nutrition, IHU-ICAN, ANR-10-IAHU-05) at Paris-Sorbonne Université, and HistIM, GENOM’IC and CYBIO core facilities at Institut Cochin. This work was supported by grants from the National Agency for Research (ANR) (ANR-GutBarrIR), the Foundation for Medical Research (FRM), the French Diabetes Society (SFD) and the European Foundation for the study of Diabetes (EFSD).

---

**Figure S1:**
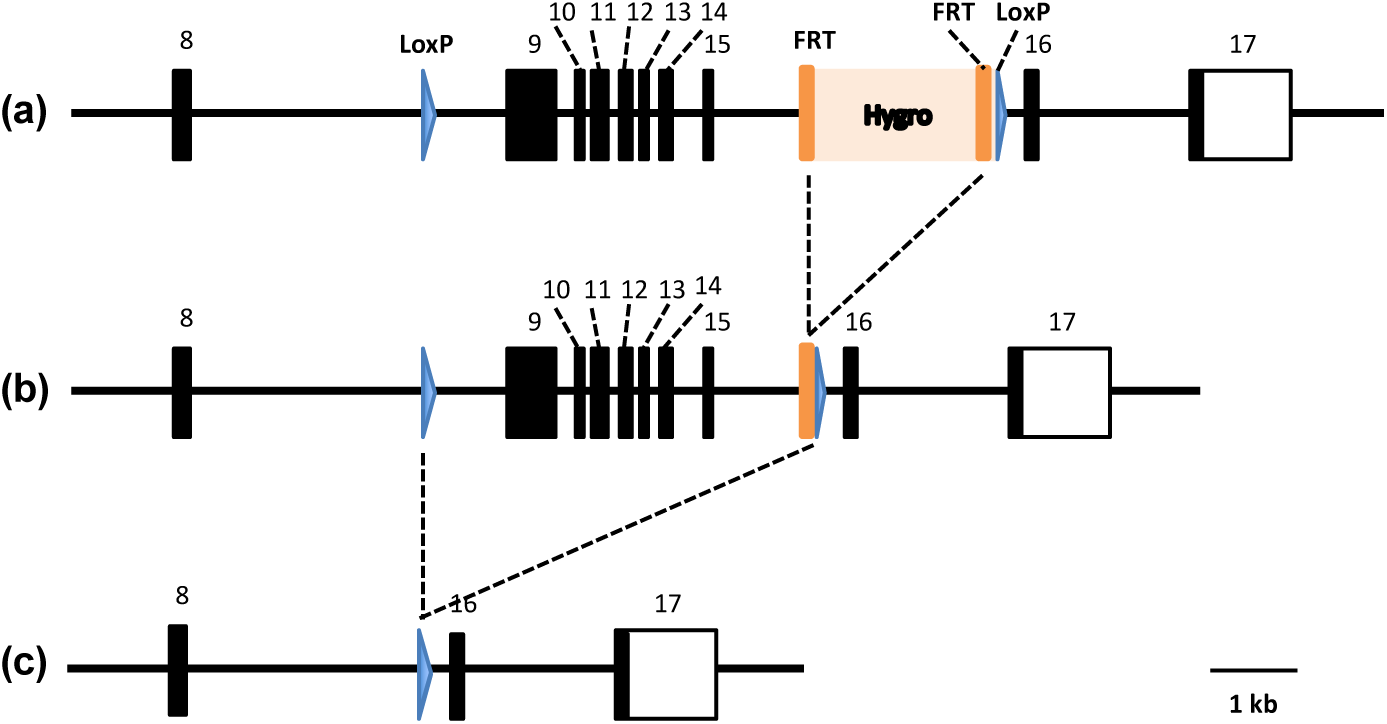
Generation of Ch^ΔGUT^mice. **(a)** Vector used to generate founder mice and containing the hygromycin cassette inserted between Exon 15 and Exon 16 of *mlxipl*, which is flanked by FRT (Flippase Recognition Target) sites. Two loxP sites flank exon 9 and hygromycin cassette. **(b)** Founder mice were crossed with mice that express the Flippase recombinase to remove the hygromycin cassette generating *mlxipl*^fl/fl^ mice. **(c)** *mlxipl*^fl/fl^ mice were then crossed with a *Villin-Cre^ERT2^* mouse line to generate *mlxipl*^fl/fl^ × *Villin-Cre^ERT2^* mice, in which the Cre recombinase was activated by tamoxifen gavage (1 mg/mouse) for 5 consecutive days to induce a specific *mlxipl* deletion in intestinal epithelial cells (Ch^ΔGUT^ mice).

**Figure S2:**
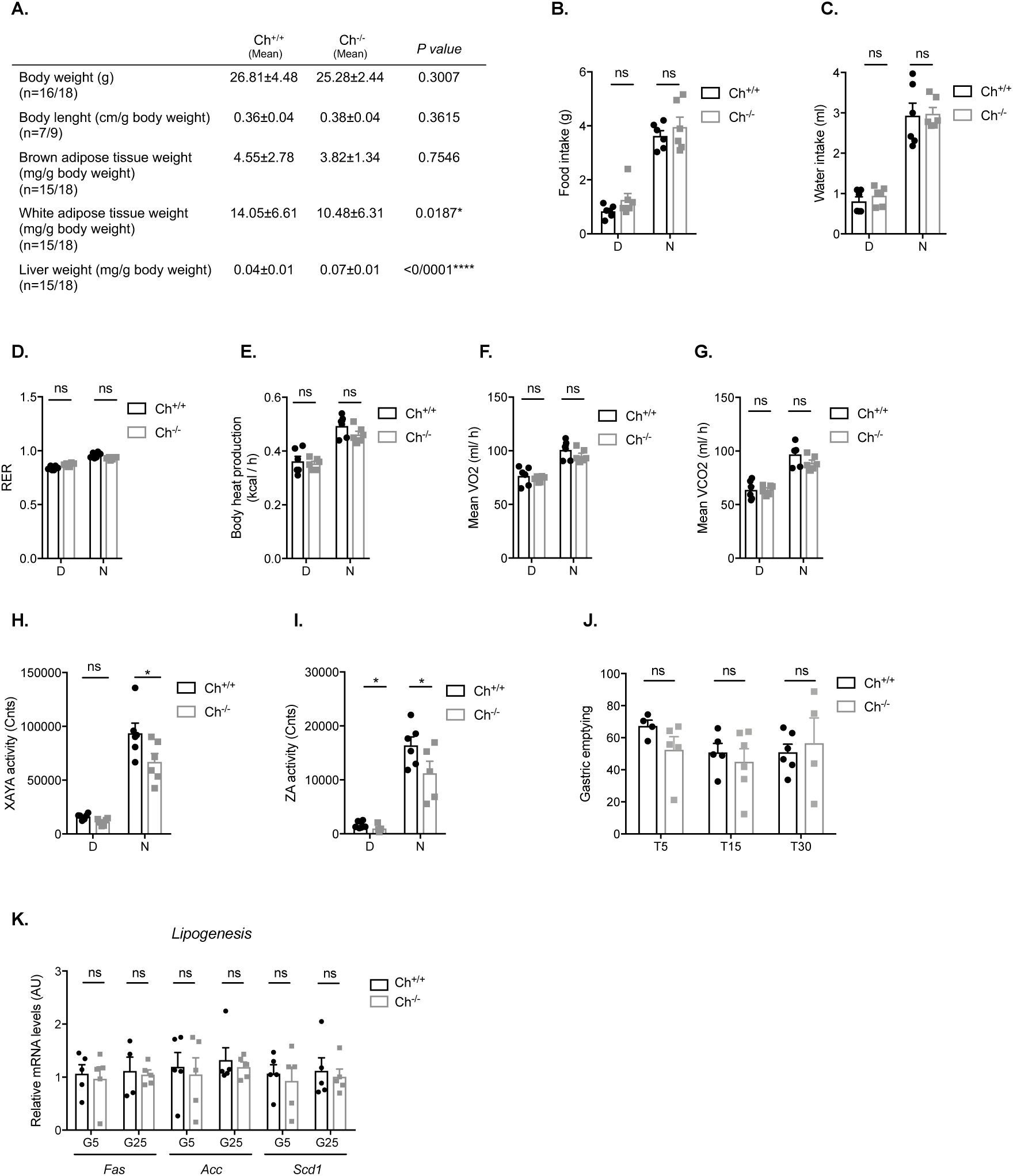
Phenotypic characteristics of ChREBP^-*/*-^ mice. **(A)** Relative tissue weights (normalized to body weight) **of** random fed 12-20-week-old Ch^+/+^ and Ch^-*/*-^ mice (n=7-18). **(B)** Food intake, **(C)** water intake, **(D)** respiratory exchange ratio (RER), **(E)** body heat production, **(F)** mean oxygen consumption (VO2), **(G)** mean carbon dioxide production (VCO2), **(H)** Horizontal activity (XAYA), and **(I)** vertical activity (ZA), measurements acquired during the day (“D”) and the night (“N”) for both Ch^+/+^ and Ch^-*/*-^ mice (n = 6)**. (J)** Gastric emptying rate assayed by the phenol red method and expressed as % of T0 (n = 4-6). **(K)** Relative mRNA levels of *fas*, *acc and scd-1* (normalized to *TBP*) in gut epithelial cells isolated from 10- to 12- weeks-old male Ch^+/+^ and Ch^-*/*-^ mice after overnight fasting. Intestinal segments from Upper Small Intestine were treated *ex vivo* for 4h with 5mM or 25mM glucose before epithelial cell isolation (n = 5). Values are means ± SEM. *\*P < 0.05, ****P < 0.0001*; compared Ch^+/+^ *vs* Ch^-*/*-^ mice (Mann Whitney U test).

**Figure S3:**
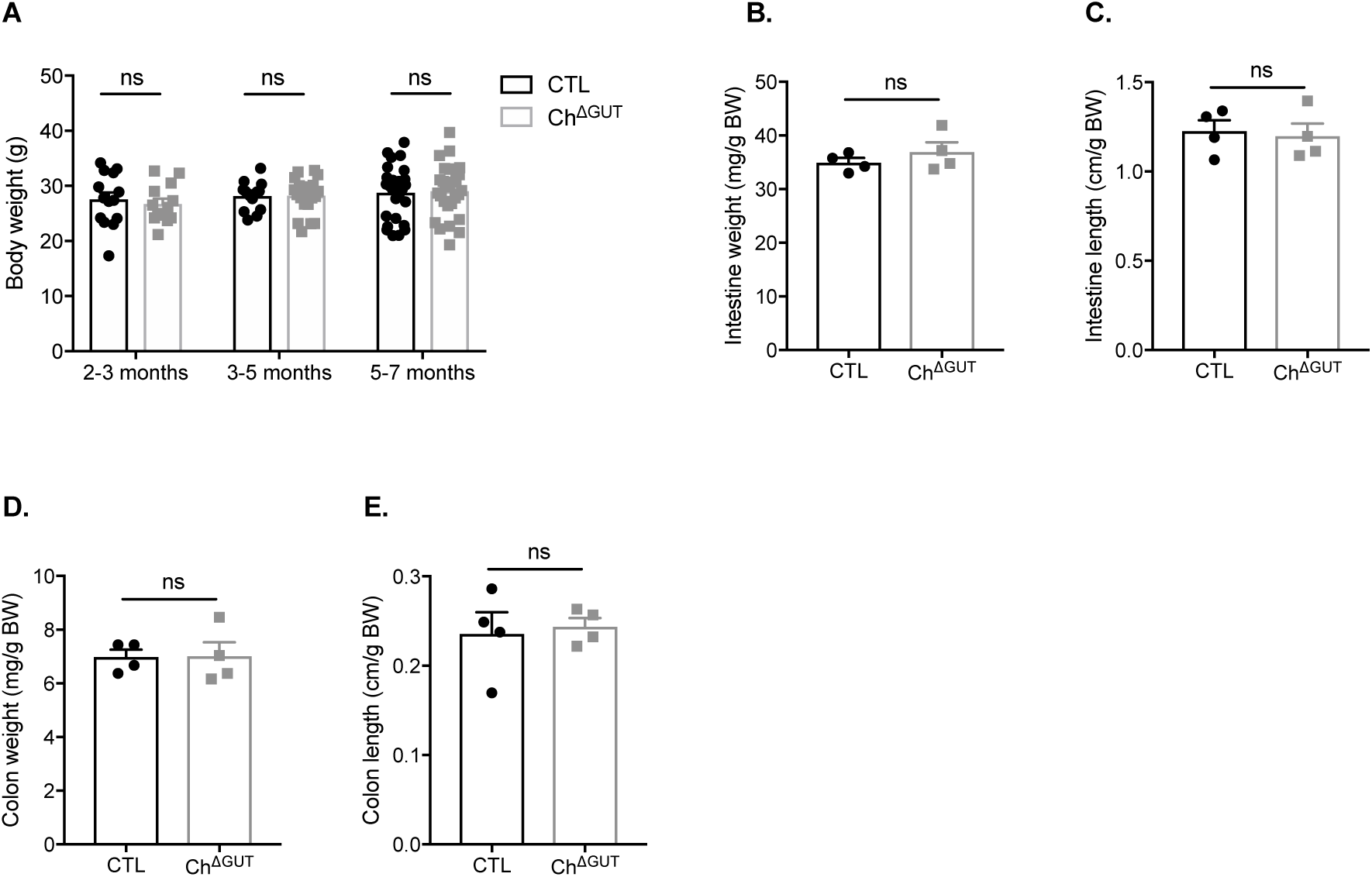
Phenotype characteristics of Ch^ΔGUT^mice. **(A)** Body weight of fasted 16-week-old CTL and Ch^ΔGUT^ mice (n = 4). **(B-E)** Relative weight and length of the small intestine and the colon of fasted 16-week-old CTL and Ch^ΔGUT^ mice (normalized to body weight) (n = 4). Values are means ± SEM. *ns*; compared CTL *vs* Ch^ΔGUT^ mice (Mann Whitney U test).

**Figure S4:**
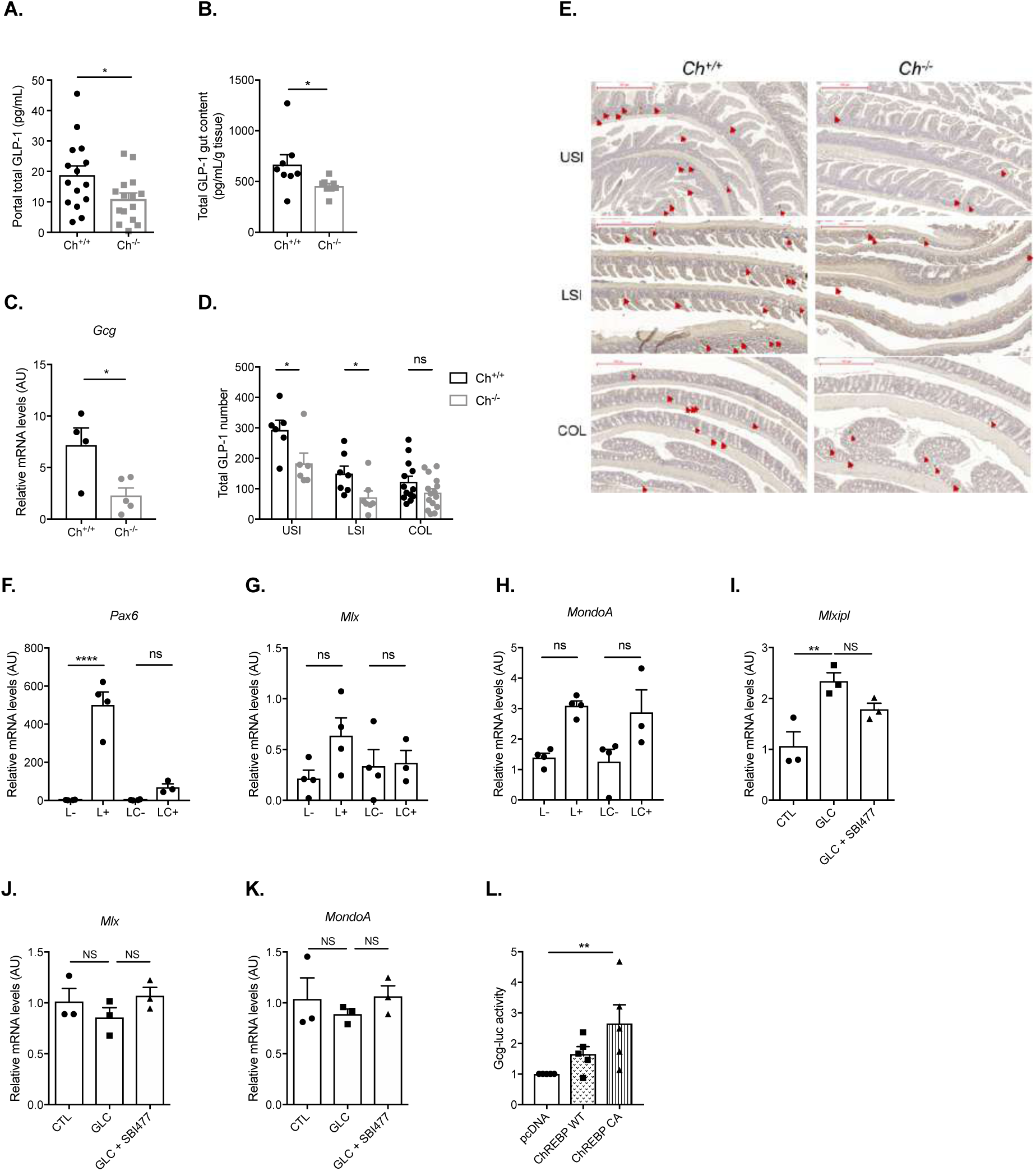
GLP-1 production is reduced upon whole body ChREBP deficiency. **(A-B)** Total GLP-1 concentrations in the portal blood (A, n = 15) or in the upper small intestinal mucosa (B, n = 8) of 16-week-old female Ch^+/+^ and Ch^-*/*-^ mice at 15 min after an oral glucose bolus (2g/kg). **(C)** *gcg* relative mRNA levels (normalized to TBP) in gut epithelial cells isolated from 10- to 12- week-old male Ch^+/+^ and Ch^-*/*-^ mice. Segments from upper small intestine were treated *ex vivo* for 4h with 25mM glucose before epithelial cell isolation (n = 4-5). **(D)** Representative images of GLP-1 immunostaining on transversal sections of the upper small intestine (USI), lower small intestine (LSI) and the colon (COL) from Ch^+/+^ *vs* Ch^-*/*-^ mice and corresponding quantification of total GLP-1 positive L-cells number per intestinal segment (n = 7-15). **(A-D)** Values are means ± SEM. *\*P < 0.05*; compared Ch^+/+^ *vs* Ch^-*/*-^ mice (Mann Whitney U test). **(E-G)** Relative mRNA levels of *pax6, mlx* and *mondoA* (normalized to *TBP*) in GLP-1 producing epithelial cells (jejunal L+ and colonic LC+ cells) and in GLP-1 negative epithelial cells (jejunal L- and colonic LC-) of 10- to 12-week-old male GLU-Venus mice (n = 3-4). Values represent means ± SEM. *ns, ****P < 0.0001*; compared L^-^ *vs* L^+^ and LC^-^ *vs* LC^+^ (one-way ANOVA). **(H)** *mlxipl* relative mRNA levels (normalized to *TBP*) in jejunal L+ and colonic LC+ cells of 10- to 12-week-old male GLU-Venus and GLU-Venus Ch^-*/*-^ mice. Values are means ± SEM (n = 3-4). ns, *\*\*P < 0.01*; compared GLU-Venus mice *vs* GLU-Venus Ch^-/-^ mice (two-way ANOVA followed by Bonferroni correction for multiple comparisons). **(I-K)** Relative mRNA levels of *mlxipl*, *mlx* and *mondoA* (normalized to *TBP*) in GLUTag cells incubated 24h with 5mM or 25mM glucose or SBI477 (10µM) in 25mM glucose. Values are means ± SEM (n = 3). *ns, **P < 0.01;* compared high glucose condition to the untreated condition (Vehicle), and SBI477 compared to high glucose condition (Mann Whitney U test)**. (F)** Relative *Gcg*-luciferase reporter activity in GLUTag cell line transfected with a plasmid encoding a wild type (ChREBP-WT) or a constitutively active form of ChREBP (ChREBP-CA) and, incubated with 5mM glucose (n = 5). The Renilla luciferase reporter pRL-CMV was used as internal control. Dual luciferase reporter assays were performed 24 h post transfection.

**Figure S5:**
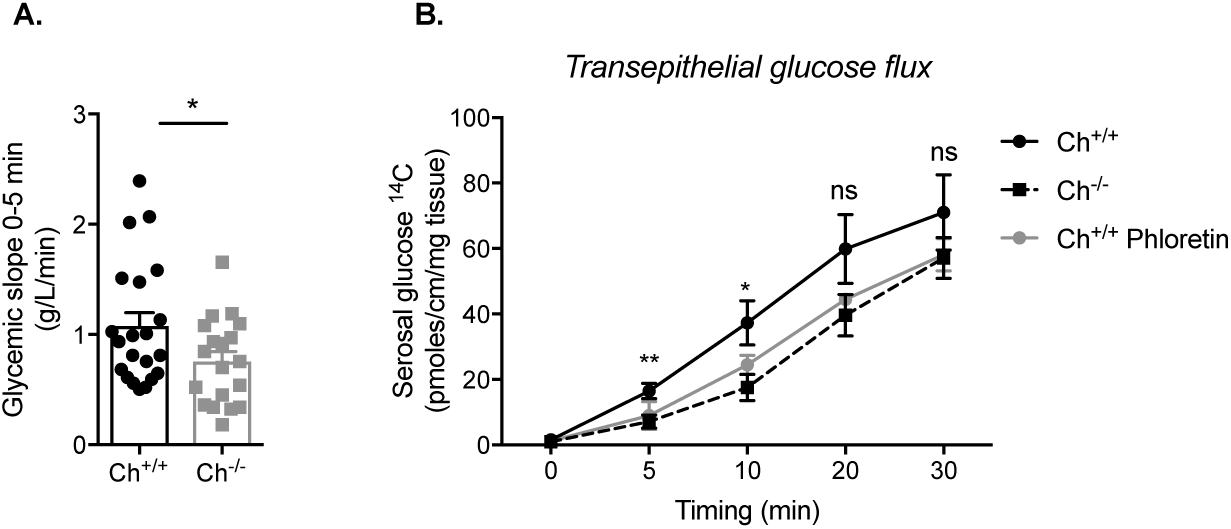
Transepithelial intestinal glucose transport is dampened upon whole body ChREBP deficiency. **(A)** Glucose absorption index as measured by the glycemic slope between 0 and 5 min after oral glucose gavage (4g/kg) in 10- to 12-week-old male Ch^+/+^ and Ch^-*/*-^ mice (n = 19-21). **(B)** Transepithelial glucose flux in jejunal loops from 10- to 12-week-old male Ch^+/+^ mice incubated with or without phloretin and from Ch^-*/*-^ mice (n = 5-6). **(A-B)** Values are means ± SEM. *ns*, *\*P < 0.05, **P < 0.01*; compared Ch^+/+^ *vs* Ch^-*/*-^ mice. (Mann Whitney U test).

**Figure S6:**
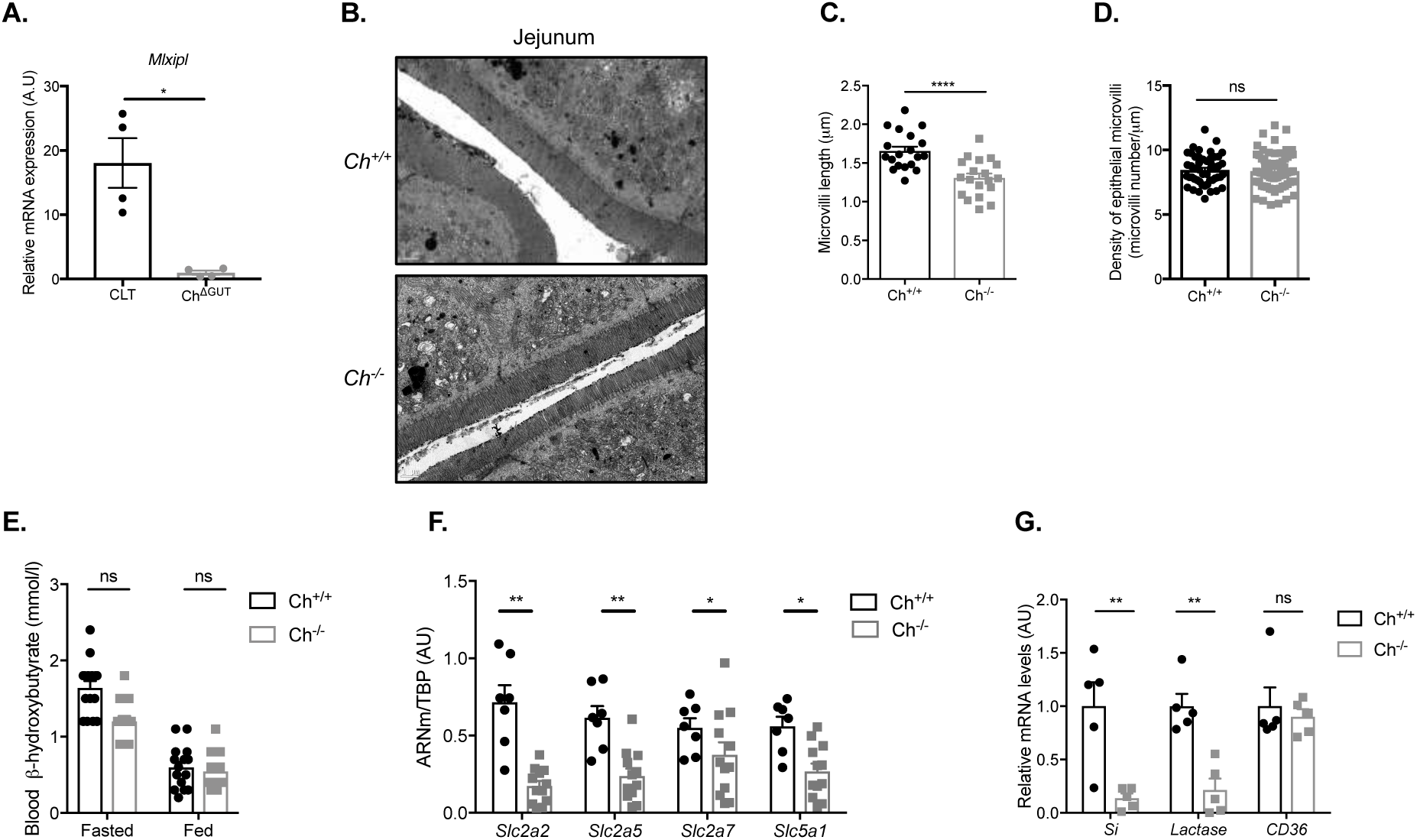
Decreased sugar digestion and absorption markers upon whole body ChREBP deficiency. **(A)** Relative *mlxipl* mRNA levels (normalized against *TBP*) in jejunal epithelial cells of 16-week-old male CTL and Ch^ΔGUT^ mice in response to an overnight oral 20%-glucose challenge. Values are means ± SEM (n = 4). *\*P < 0.05*; compared CTL *vs* Ch^ΔGUT^ mice (Mann Whitney U test). **(B)** Representative transmission electron photomicrographs of the jejunal brush border from 10- to 12-week-old random fed male Ch^+/+^ and Ch^-*/*-^ mice (Magnification ×2000). Scale bar = 1µm. **(C-D)** Microvilli length and density in jejunum of 16-week-old male CTL and Ch^ΔGUT^ mice (n = 4). **(E)** β-hydroxybutyrate plasma levels in 10- to 12-week-old fasted or fed male Ch^+/+^ and Ch^-*/*-^ mice (n = 15). **(F-G)** Relative mRNA levels of hexose transporters and disaccharidases (normalized against *TBP*) in jejunal epithelial cells from 10- to 12-week-old male Ch^+/+^ and Ch^-*/*-^ mice (n = 5-12). **(C-G)** Values are means ± SEM. *ns, *P < 0.05, **P < 0.01, ****P < 0.0001*; compared Ch^+/+^ *vs* Ch^-*/*-^ mice (Manne Whitney U test).

**Figure S7:**
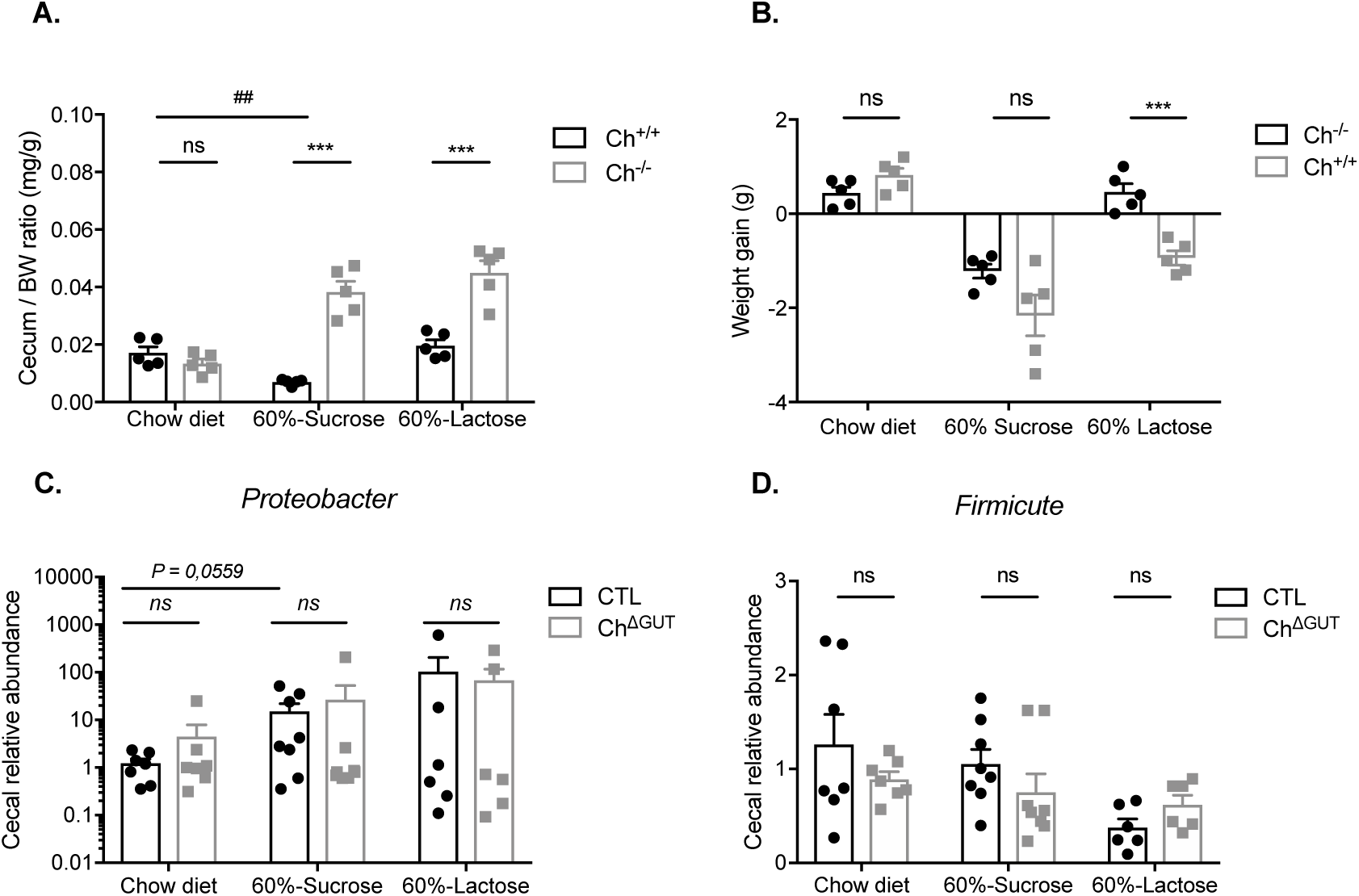
Total ChREBP deficiency induces early intolerance to high sucrose and lactose diets. **(A-B)** Relative caecum weight (normalized to body weight) and body weight gain in 10- to 12-week-old male Ch^+/+^ and Ch^-*/*-^ mice fed either with chow, 60%-sucrose or 60%-lactose diet for four days (D4) (n = 5). Values are means ± SEM. *ns, ***P < 0.001*; compared Ch^+/+^ *vs* Ch^-*/*-^ mice (Manne Whitney U test). Values are means ± SEM. *ns, ***P < 0.001*; compared Ch^+/+^ *vs* Ch^-*/*-^ mice (Mann Whitney U test).

**Figure S8:**
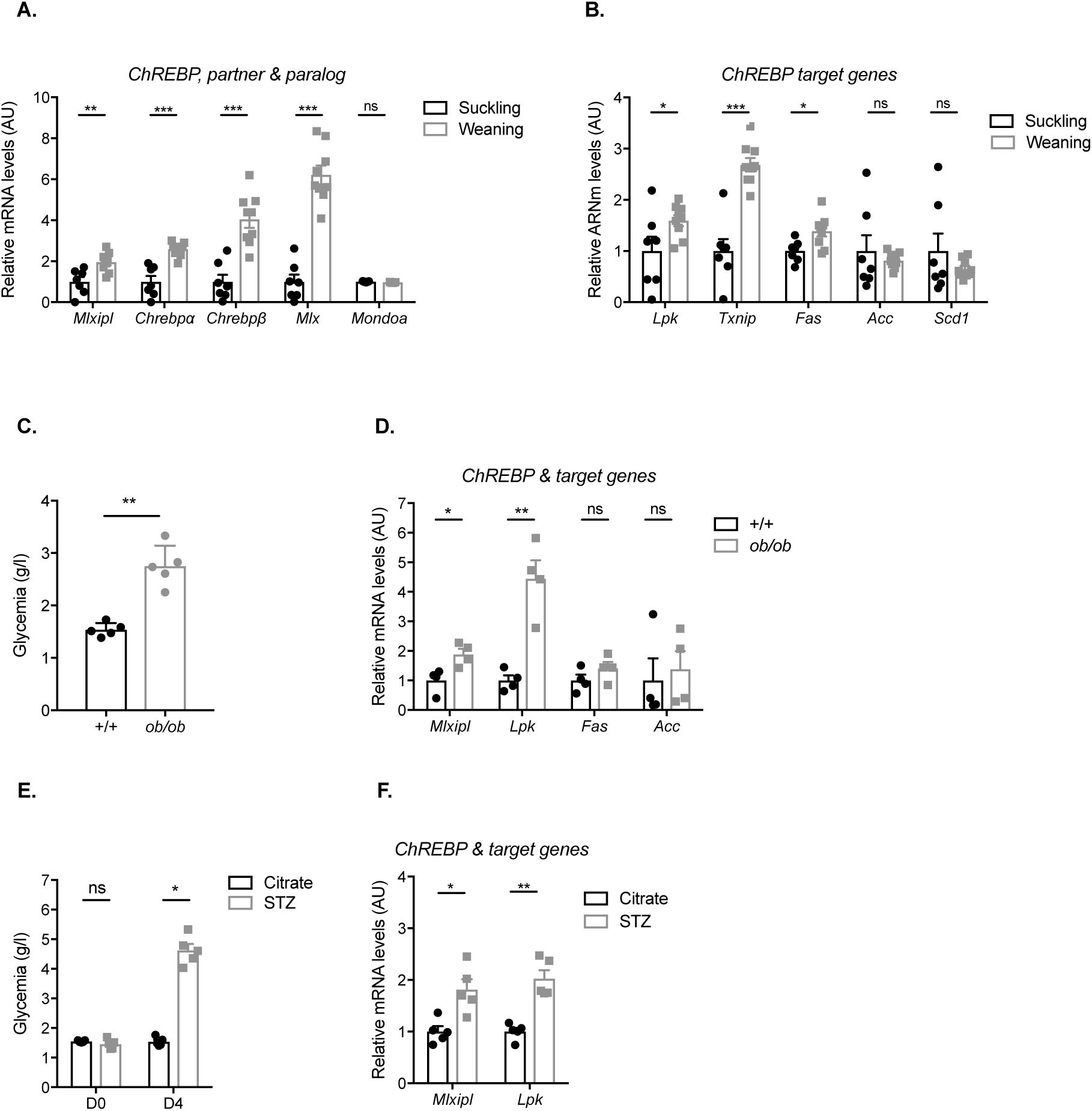
Characterization of ChREBP expression and activity upon weaning or hyperglycemic conditions. **(A-B)** Relative mRNA levels (normalized to *TBP*) in small intestinal epithelial cells of 8 (suckling) or 28 day-old (weaning) wild type mice. Values are means ± SEM (n = 7-9). *ns, *P < 0.05, **P < 0.01, ***P < 0.001*; compared suckling *vs* weaning mice (Mann Whitney U test). **(C)** Plasma glucose and **(D)** relative mRNA levels (normalized to *TBP*) in jejunal epithelial cells of 12-week-old male wild type (+/+) and obese (*ob/ob*) mice. Values are means ± SEM (n = 4). *ns, *P < 0.05, **P < 0.01*; compared wild type *vs ob/ob* mice (Mann Whitney U test). **(E)** Plasma glucose and **(F)** relative mRNA levels (normalized to *TBP*) in jejunal epithelial cells from 10-week-old male mice treated for 5 days with buffer (Citrate) or streptozotocin (STZ). Values are means ± SEM (n = 5). *ns*, *\*P < 0.05;* compared PBS *vs* STZ mice (Mann Whitney U test). Values are means ± SEM (n = 4). *\*P < 0.05, **P < 0.01*; compared PBS *vs* STZ mice (Mann Whitney U test).

